# Mitochondrial genomes in *Perkinsus* decode conserved frameshifts in all genes

**DOI:** 10.1101/2022.02.17.480897

**Authors:** Sebastian G. Gornik, Victor Flores, Franziska Reinhardt, Lieselotte Erber, Olga Douvropoulou, Imen Lassadi, Elin Einarsson, Mario Mörl, Anna Git, Peter F. Stadler, Arnab Pain, Ross F. Waller

## Abstract

Mitochondrial genomes of apicomplexans, dinoflagellates and chrompodellids, that collectively make up the Myzozoa, are uncommonly reduced in coding capacity and display divergent gene configuration and expression mechanisms. They encode only three proteins — COB, COX1, COX3 — contain rRNAs fragmented to ∼100-200 base pair elements, and employ extensive recombination, RNA trans-splicing, and RNA-editing for genome maintenance and expression. The early-diverging Perkinsozoa is the final major myzozoan lineage whose mitochondrial genomes remain poorly characterized. Previous reports of *Perkinsus cox1* and *cob* partial gene sequences have indicated independent acquisition of non-canonical features, namely the occurrence of multiple frameshifts in both genes. To determine ancestral myzozoan mitochondrial genome features, as well as any novel ones in Perkinsozoa, we sequenced and assembled four *Perkinsus* species mitochondrial genomes. These data show a simple ancestral genome with the common reduced coding capacity, but one already prone to rearrangement. Moreover, we identified 75 frameshifts across the four species that are present in all genes, that are highly conserved in gene location, and that occur as four distinct types. A decoding mechanism apparently employs unused codons at the frameshift sites that advance translation either +1 or +2 frames to the next used codon. The locations of the frameshifts are seemingly positioned to regulate protein folding of the nascent protein as it emerges from the ribosome. COX3 is distinct in containing only one frameshift and showing strong selection against residues that are otherwise frequently encoded at the frameshift positions in COX3 and COB. All genes also lack cysteine codons implying a further constraint on these genomes with reduction to only 19 different amino acids. Furthermore, mitochondrion-encoded rRNA fragment complements are incomplete in *Perkinsus* spp. but some are found in the nuclear DNA, suggesting these may be imported into the organelle as for tRNAs. *Perkinsus* demonstrates additional remarkable trajectories of organelle genome evolution including pervasive integration of frameshift translation into genome expression.

## INTRODUCTION

Mitochondria are morphologically distinctive double-membrane organelles best known for their role in ATP synthesis in eukaryotic cells via oxidative phosphorylation (Saraste 1999; Gray et al. 2001; Burger et al. 2003; Wang and Youle 2009; Flegontov et al. 2015; Roger et al. 2017). Mitochondria also contribute to numerous other important cellular functions including iron-sulfur (Fe/S) cluster biogenesis, anabolic metabolism (e.g. heme biosynthesis) and apoptosis (Saraste 1999; Gray et al. 2001; Burger et al. 2003; Wang and Youle 2009; Flegontov et al. 2015; Roger et al. 2017). All extant mitochondria are derived from an ancient endosymbiosis of an alpha-proteobacterium within the eukaryotic common ancestor, and most have retained some form of the original prokaryotic genome (Roger et al. 2017). However, despite their common origin and generally conserved functions, mitochondrial genomes (mtDNAs) display a remarkable diversity of states (Gray et al. 2001; Burger et al. 2003; Smith and Keeling 2015; Gagat et al. 2017; Roger et al. 2017; Berná et al. 2021).

All mtDNAs are vastly reduced compared to the progenitor prokaryotic genomes. Retained genes typically code for components of the electron transport chain (complexes I, III, IV and V) and the mitochondrial translation machinery, notably transfer RNAs (tRNAs) and ribosomal RNAs (rRNAs) (Lang et al. 1997; Roger et al. 2017). The protein-coding capacity of mtDNAs ranges from near 100 proteins in jakobid flagellates (Lang et al. 1997; Burger et al. 2003; Flegontov et al. 2015) to a mere two in the chrompodellid *Chromera velia*, with the typical number across eukaryotes being 40–50 genes (Gray et al. 2001; Burger et al. 2003; Flegontov et al. 2015; Roger et al. 2017). This surprisingly large divergence across extant mtDNAs is the consequence of both organelle function loss and ongoing endosymbiotic gene transfer to the nucleus occurring throughout eukaryotic diversification (Gray et al. 2001; Burger et al. 2003; Herrmann 2003; Roger et al. 2017). The relocated genes are translated in the cytosol and the corresponding proteins imported back into the organelle (Herrmann 2003; Maguire and Richards 2014; Roger et al. 2017). In some instances mitochondria are so derived, reduced and specialized (e.g. hydrogenosomes and mitosomes) that they lack the electron transport chain components and have lost their mtDNAs completely (Maguire and Richards 2014; Roger et al. 2017; Berná et al. 2021). The architecture of the remaining organelle genomes are primarily linear, single chromosomes, which often appear circular in sequence assemblies due to various stabilizing inverted end-structures such as terminal inverted repeats (TIRs) that cause false in-silico circularization (Cavalier-Smith 2018; Berná et al. 2021).

The Myzozoa, which comprise dinoflagellates, apicomplexans, chrompodellids and perkinsids (Flegontov et al. 2015; Roger et al. 2017; Cavalier-Smith 2018), represent some of the most reduced and divergent mtDNAs known to date (Flegontov et al. 2015; Roger et al. 2017). They encode the smallest number of proteins for any organelle genome — COB, COX1, COX3 — with the gene for COB apparently completely lost in the chrompodellid *Chromera velia* (Waller and Jackson 2009; Flegontov et al. 2015; Roger et al. 2017; Berná et al. 2021). Moreover, rRNAs in myzozoan mtDNAs are highly fragmented with no evidence of reassembly at the RNA level by splicing, and they entirely lack 5S rRNA (Waller and Jackson 2009; Feagin et al. 2012; Berná et al. 2021). The architecture of myzozoan mtDNAs is multiform and complex. For example, most apicomplexan mtDNAs that have been characterized to date have monomeric compacted linear forms, as small as 6 kb in *Plasmodium* (Feagin et al. 2012; Berná et al. 2021; Namasivayam et al. 2021). *Toxoplasma gondii*, however, has diverged from other apicomplexans with highly expanded, fragmented, but modularized mtDNAs (Flegontov et al. 2015; Namasivayam et al. 2021). Within the apicomplexan sister group, the chrompodellids, expansion of mtDNAs is also seen as heterogeneous, linear, duplicated and fragmented molecules (Slamovits et al. 2007; Waller and Jackson 2009; Jackson et al. 2012; Flegontov et al. 2015), and a similar pattern is found yet again in dinoflagellates surveyed to date (Jackson et al. 2007; Slamovits et al. 2007; Nash et al. 2008; Waller and Jackson 2009; Jackson et al. 2012; Gagat et al. 2017). This distribution of both simple and complex mtDNAs throughout Myzozoa confuses understanding of the ancestral state of the myzozoan mtDNA: was it monomeric or heterogeneous? Furthermore, dinoflagellate mtDNAs possess additional features and traits, namely: 1) extensive substitutional RNA editing, 2) trans-splicing of *cox3* transcripts, 3) use of alternative start codons (also found in apicomplexans); and 4) general loss of encoded stop codons (Jackson et al. 2007; Nash et al. 2008; Waller and Jackson 2009; Jackson et al. 2012; Gagat et al. 2017; Janouskovec et al. 2017). Of particular note, *cob* and *cox1* mRNAs are poly-adenylated immediately after the region encoding the conserved C-terminus, presumably resulting in a short read-through poly-lysine tail. However, the molecular evolutionary behavior of *cox*3 differs in that poly-adenylation consistently creates a UAA in-frame stop codon (Waller and Jackson 2009; Jackson et al. 2012).

The Perkinsozoa represents the deepest-branching lineage of the dinozoan clade diverging relatively close to the split with the sister apicomplexan/chrompodellid clade (Masuda et al. 2010; Zhang et al. 2011; Janouskovec et al. 2017). Thus, Perkinsozoa represents an early branching point of myzozoan diversity. *Perkinsus* spp. are the best studied representatives of Perkinsozoa owing to their importance as marine parasites and pathogens of commercial important shellfish (Villalba et al. 2007; Choi and Park 2010; Smolowitz 2013). To date, only partial coding sequence for two mitochondrial genes has been reported from *Perkinsus* spp. — *cob* and *cox1* — yet, these already indicate some further novelty in myzozoan mtDNAs (Masuda et al. 2010; Zhang et al. 2011; Bogema et al. 2021). These sequences indicate the use the canonical UGA stop codon as an alternative code for tryptophan, which is a trait seen in other mitochondria including ciliates although not known from other myzozoans. More radical is the observation that both coding sequences contain multiple frameshifts that are apparently not resolved by editing at the mRNA stage. These frameshifts correlate with conserved glycine and proline residues, and the nucleotide sequence at these sites indicates consensus sequences for the glycine-encoded frameshifts (AGGY or TMGGY) and proline-encoded frameshifts (CCCCT). To date, attempts to identify a *cox3* coding sequence have been unsuccessful and it has been speculated as being lost (Koren et al. 2017; Bogema et al. 2021).

While a tantalizing glimpse of the mtDNA in *Perkinsus* has been provided, there are many unanswered questions for this key lineage: What is the form of their mtDNA, monomeric or heterologous? What is its complete coding capacity of *Pekinsus* mitochondria? How are the frameshifts decoded? Can *Perkinsus* inform on the likely ancestral state of the myzozoan mtDNA? To address these questions, we sequenced and assembled the mtDNA from four *Perkinsus* taxa: *P. atlanticus, P. olseni, P. chesapeaki* and *P. marinus*, defining the structures, coding capacity, and remarkable use of frameshifts in yet another form of divergent myzozoan mtDNAs.

## RESULTS

### *Perkinsus* mtDNAs comprise unique, circular-mapping molecules

To determine the form of *Perkinsus* mitochondrial genomes (mtDNAs) and its evolutionary stability we selected four taxa for good representation of this genus. A *Perkinsus* molecular phylogeny shows some persistent ambiguity in the taxonomy of this group, nevertheless the isolates used in this study all have distinct nuclear genomic sequence (data not shown) and represent diversity across this genus: *P. atlanticus* (ATCC 50984), *P. olseni* (ATCC PRA-205), *P. chesapeaki* (ATCC 50807), and *P. marinus* (ATCC 50983) (Figure 1A). Mitochondrial genomes were assembled from short-read and long-read total DNA sequence data, and all assemble mtDNAs were circular-mapping single molecules with the exception of the *P. chesapeaki* mtDNAs that comprised two separate molecules (Figure 1B). The sizes of total mitochondrial genomes ranged from 95,104 bp (*P. atlanticus*) to 40,900 bp (*P. marinus*). All *Perkinsus* mtDNAs were strongly AT-skewed ranging from 82.2% (*P. marinus*) to 86.3% AT content (*P. chesapeaki*). Mitochondrial genome structural comparisons indicated little synteny between the four taxa (Figure 1B), with shared sequence elements limited to protein- and rRNA-coding sequences (see sections below). Greatest conservation of sequence was seen between the more closely related *P. atlanticus* and *P. olseni* (Supplementary Figure S1), however, even here there is evidence of considerable genome rearrangement and unique sequence.

**Figure 1.**
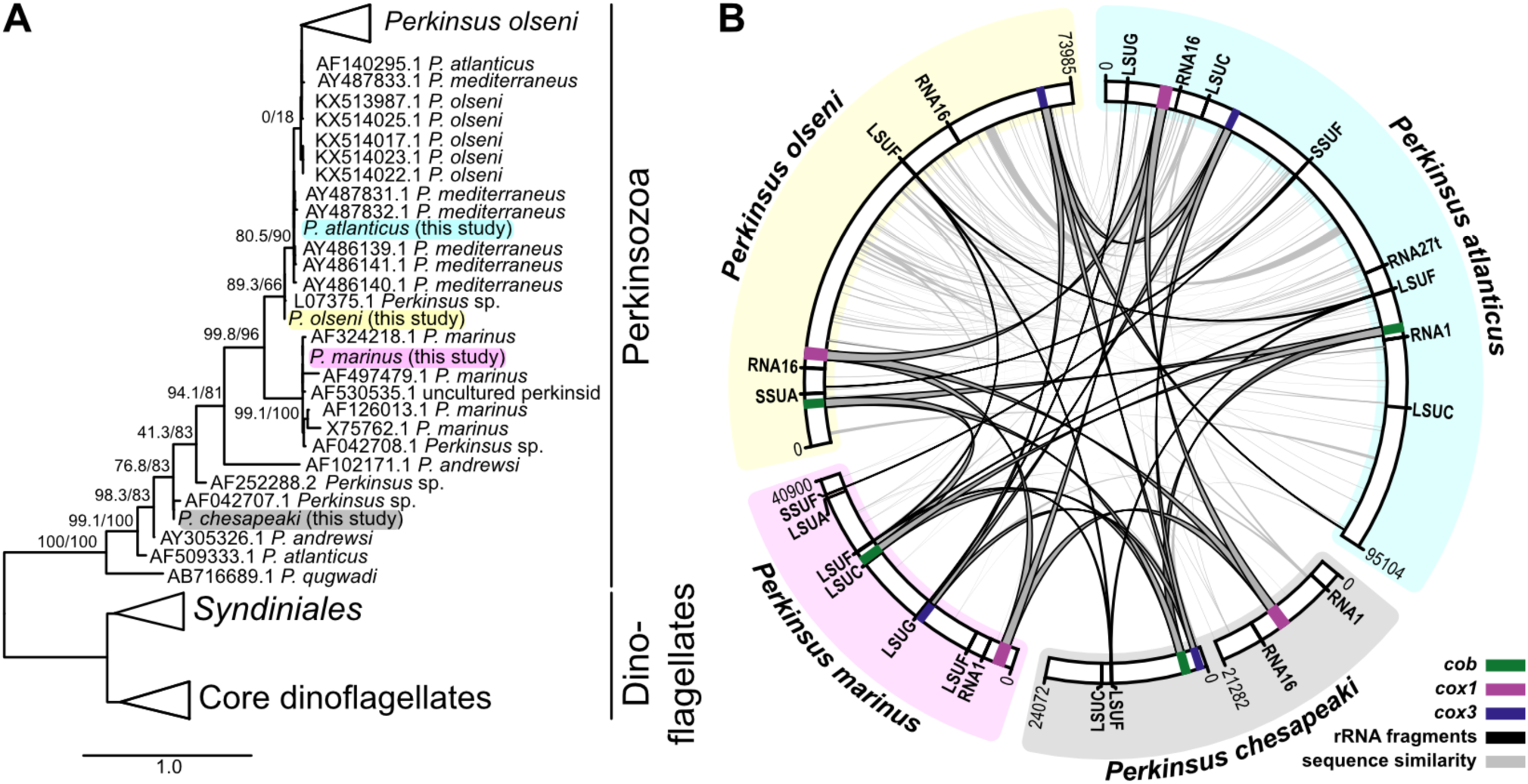
Four *Perkinsus* representatives contain simple mtDNAs with limited sequence conservationn. ***A***. An 18S SSU maximum likelihood phylogeny shows the relationship of the four taxa used in this study. Main branch support are: SH-like approximate likelihood ratio test (SH-aLRT) and Ultrafast bootstrap (UFBoot). Full phylogeny available in Supplemental Figure 1. ***B***. Overview of the mitochondrial genomes (mtDNAs) of four *Perkinsus* species. The outer ring segments represent the mtDNA, the coding regions are shown as colored bands and the predicted rRNA fragments are shown as black lines. The arcs connecting the segments of each mtDNA represent sequence similarity (syntenic) regions (reciprocal BLASTn hits with an e-value less or equal to 1×10^−3^). Note: the *P. chesapeaki* mtDNA comprises two molecules. Fully annotated genome sequences provided in Genbank format in Supplementary File 2.

### Protein coding content of the *Perkinsus* mtDNA genomes

Comparison of the four *Perkinsus* spp. mtDNAs showed conservation of only three putative protein-coding genes (Figure 1B). This is consistent with the highly reduced coding capacity seen in all Myzozoan mtDNAs containing only genes for cytochrome B (*cob*) and cytochrome C oxidase subunits 1 and 3 (*cox1* and *cox3*, respectively). No other putative protein-coding sequences were apparent in the *Perkinsus* mtDNAs. *Perkinsus cox1* and *cob* genes had been previously reported, but sequence for *cox3* had not, and this was suggested to have been lost (Masuda et al. 2010; Zhang et al. 2011; Jackson et al. 2012; Bogema et al. 2021). Indeed, searching our mtDNAs with diverse *cox3* sequences failed to identify a putative *Perkinsus cox3*. However, the predicted translation of the third common *Perkinsus* mtDNA coding sequence contains the conserved superfamily domain SSF81452 (Cytochrome c oxidase subunit III-like; evalue: 8.76×10^−8^) and IPR domain IPR013833/G3DSA:1.20.120.80 (Cytochrome c oxidase, subunit III, 4-helical bundle; e-value: 1.4×10^−6^) consistent with this ORF representing a divergent *cox3*. Protein structural modelling of the *Perkinsus* COX3 sequences (see below) further supports that these sequences represent bona-fide *cox3*.

Alignments of the four *Perkinsus* taxa for the three mitochondrial genes shows very high nucleotide identity within the conserved predicted protein coding sequence, and then dramatic loss of conservation in the immediate flanking sequence (Figures 2, S2-4). No potential ATG start codons are seen at the 5’ regions of these sequences suggesting a likely alternative initiator codon(s). Conserved asparagine (N) and phenylalanine (F) codons occur at the 5’ ends of all three genes, and either of them might be the N-terminal translation initiator of these genes. *Perkinsus* taxa are predicted to use the terminator codon TGA as an alternative tryptophan codon (Masuda et al. 2010; Zhang et al. 2011), and the use of this codon is highly conserved across the four species in all three mitochondrial genes including *cox3* (Figure 3). At the *cox3* 3’ end, the high sequence identity seen for the four species ends with a conserved TAA, suggesting that this might serve as a translation termination signal (Figure 2). The *cox1* genes, however, lack a common putative stop codon, with *P. olseni* only encoding a canonical stop codon more than 100 nucleotides further downstream. The *cob* genes similarly lack a conserved stop codon at the 3’ end of sequence conservation. The *cob* genes, however, do have an internal conserved TAA (Figure S3) that was presumed to terminate translation by Zhang et al. (2011). Strong sequence identity beyond this codon shared amongst the four *Perkinsus* taxa (Figure 2), and conservation of the corresponding predicted protein with other COBs, strongly suggests that this TAA is skipped as a terminator, and we discuss a likely explanation for this below. In the absence of clear knowledge of start and stop codons used in *Perkinsus* mtDNAs we use the predicted translations of the conserved sequences for the rest of this report (see Figures S5-7).

**Figure 2.**
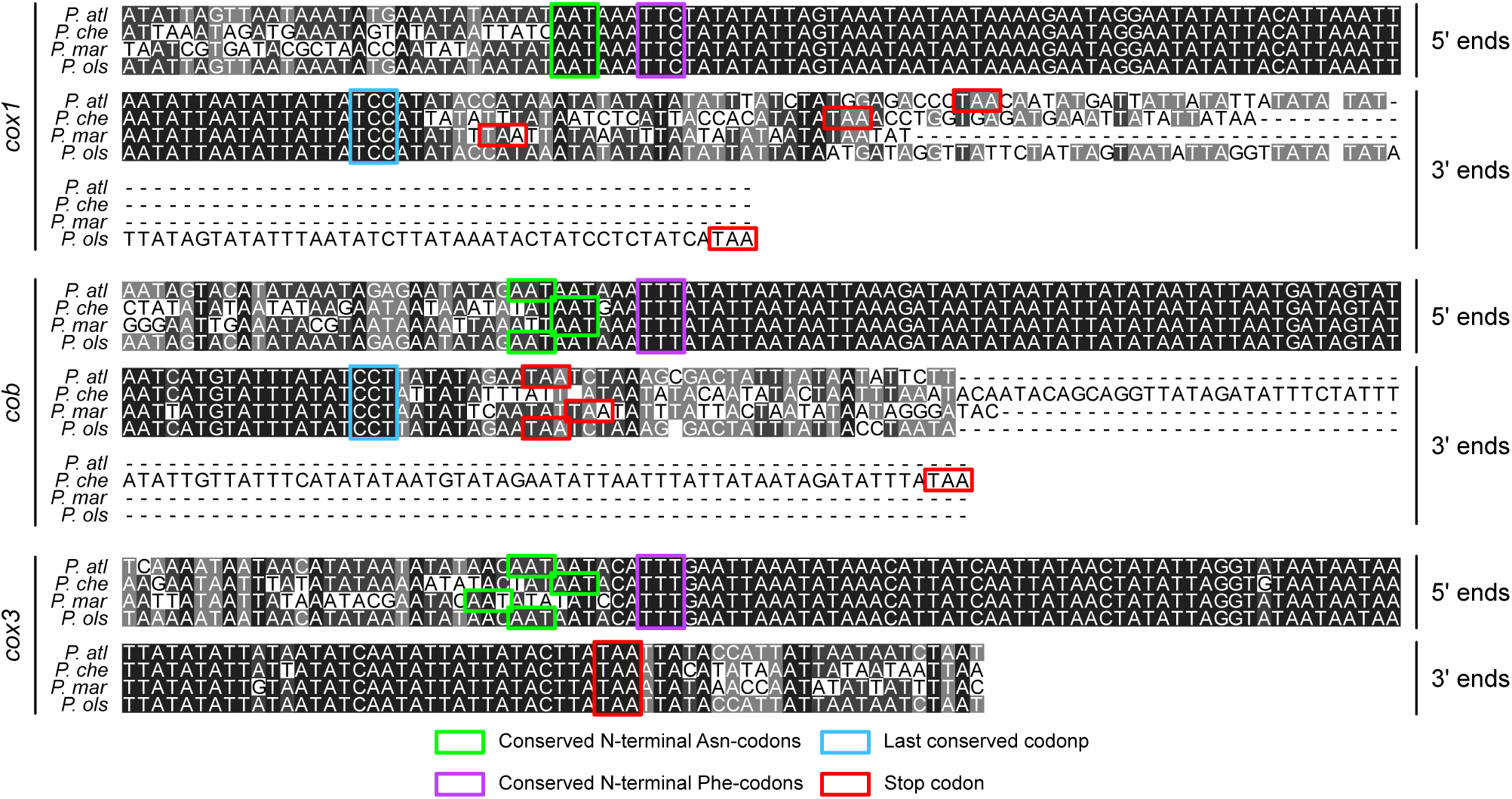
Nucleotide sequence alignments of the 5’ and 3’ ends of cox1, cob and cox3. Conserved residues as possible initiator codons, and putative terminator TAA codons, are indicated. P. alt, P. atlanticus; P. che, P. chesapeaki; P. mar, P. marinus; P. ols, P. olseni.

**Figure 3.**
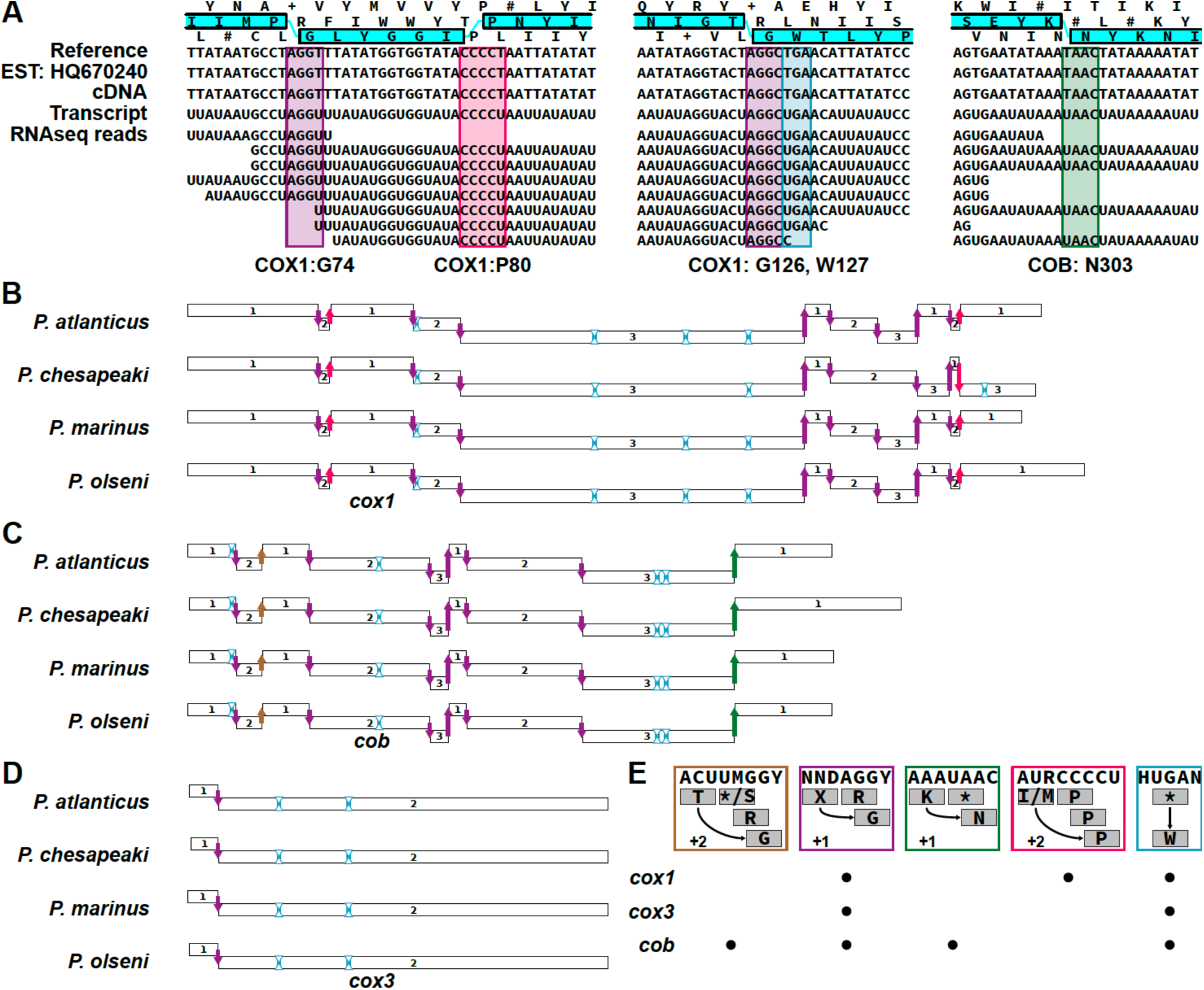
Conservation of frameshift positions in *Perkinsus* spp *cox1, cob* and *cox3*. ***A***. Representative frameshifts occurring at COX1 glycine (G) residue positions (purple boxes), COX1 proline (P) residue positions (pink box), and COB asparagine (N) residue positions (green box). The use of an alternative tryptophan (W) TGA codon is shown in the blue box. Both the frameshifts and the alternative TGA codon are present also in the RNA molecules seen by EST sequences, cDNA clones and RNA-seq data. ***B***., ***C***. and ***D***. Distribution of the G, P and N frameshifts, and the alternative W codon (blue outlined arrowheads) observed in the COX1, COB and COX3 coding regions. ***E***. Consensus sequences of the four frameshift types and alternative W codon, and comparative distribution of the alternative codes in the three genes.

### Conserved frameshifts occur in all protein genes but are strongly depleted from cox3

Previous DNA and cDNA sequencing of *P. marinus cox1* and *cob* genes identified frameshifts in these coding sequences (Masuda et al. 2010; Zhang et al. 2011; Erber et al. 2019). Only when the reading frames are shifted at multiple points is a conserved protein translation predicted (Figure S6 and S7). The positions of the predicted shifts of either +1 or +2 frames showed nucleotide sequence conservation and coincided with predicted locations for glycine residues (at sites with the consensus sequences AGGY or TMGGY) and proline residues (at sites with the consensus sequence CCCCT; Figure 3A, Figure S6 and S7). One interpretation of this is that these four- and five-nucleotide sequences serve as alternative, extended codes for glycine and prolines, although, canonical three-nucleotide codons also specify conserved glycine and proline positions elsewhere in both genes (Figure S6 and S7). The positions and sequence of the frameshifts in *cox1* and *cob* is highly conserved in *Perkinsus* (Figure 3B). Only the position of glycine G384 in *P. chesapeaki* has lost this frameshift, or never gained it. To exclude a role of RNA-editing in correcting the frameshifts, we aligned RNAseq reads, assembled transcripts and Sanger-sequenced cDNA clones to the mitochondrial genome sequences (Figure 3A for example). No differences between the genomic and transcript-derived sequences were observed confirming that no RNA-editing occurs in *Perkinsus* mitochondrial transcripts.

In addition to the frameshifts observed by Masuda et al (2010) and Zhang et al (2011), the sequence conservation of the 5’ region of *Perkinsus cob* genes downstream of the in frame TAA (Figure 2 and S4) implies a further, novel, mitochondrial frameshift. The inferred translation of the +1 frame from this TAA shows sequence conservation to the terminus of other COB proteins (Figure S7), and this predicted final reading frame is only open for four putative codons upstream of the TAA. These data suggest that a final frameshift occurs in this four-codon window before the TAA. The consensus sequences for the glycine (AGGY or TMGGY) or proline (CCCCT) frameshifts, however, do not occur in this window and we predict that an alternative frameshift occurs at this site (Figure 3C and E, and discussed below).

To query whether the newly identified *cox3* genes also contain possible frameshifts, we modeled the sequence of the most abundant frameshift (+1 frame at glycine) to search for candidate frameshift positions in the *cox3* sequences. We generated a position-specific matrix model derived from all 55 ‘AGGY’ frameshifts in *cox1* and *cob* from the four taxa, including the flanking 20 nucleotides on either side of the frameshifts. This modeling identified an enriched consensus sequence: WAWTAAWWWYT<u>AGGT</u>WTAWH AKTWATAATW (e-value: 2.7e^-149^) that, when searched against *cox3*, identified one possible frameshift in this gene in all taxa (Figure 3D). This frameshift alone is sufficient to extend the single open reading frame that spans the full conserved sequence shared amongst the four *Perkinsus* spp (Figure S7). Thus, while *cox1* and *cob* contain 10 and 8 frameshifts, respectively, *cox3* is distinct in containing only one. An interesting implication for this is a paucity of predicted glycine and proline residues in *Perkinsus* COX3 proteins. Furthermore, there are no canonical three-codon residues for glycine and proline in the *Perkinsus cox3* sequences despite COX3 proteins in other organisms containing several of those residues that are often highly conserved in position (Figure S7). These data suggest that some strong selective pressure exclusively on *Perkinsus cox3* and not *cox1* or *cob*.

### All tRNAs are imported into the mitochondria and none are predicted to decode 4/5-nucleotide codons

Given the evidence for non-canonical translation processes in the mitochondria of *Perkinsus*, we examined the tRNA pools of these cells for evidence of tRNAs that might participate in decoding the frameshifts. All known myzozoan mtDNAs lack encoded tRNAs, however, the deep branching point of Perkinsozoa within this large group allowed the possibility of retention of distinct mitochondrion-encoded tRNAs in this lineage. We scanned the mtDNAs of the four *Perkinsus* taxa with tRNAscan-SE and found no evidence of encoded tRNAs. This suggests that *Perkinsus* mitochondria are also fully dependent on importing cytosolic tRNAs. To examine the total pool of cellular tRNAs we then performed LOTTE-Seq (Long hairpin oligonucleotide based tRNA high-throughput sequencing) for both *P. marinus* and *P. olseni* from total cell RNA. This method enriches for the tRNA 3’-CCA ends (Hou 2010; Erber et al. 2019; Chan et al. 2021). We sequenced each of these samples to 100x genomic coverage and, from this, we identified 50 canonical tRNAs spanning all codon types (Supplementary Table 2). None of these *Perkinsus* tRNAs mapped to the mtDNAs suggesting that all are nucleus-encoded. The only LOTTE-Seq reads that did map to mtDNAs were AT-rich sequences that lacked predicted tRNA structure and often contained the CCA sequence in the genome but which is typically not encoded in the tRNA genes (Hou 2010; Feagin et al. 2012; Jackson et al. 2012; Flegontov et al. 2015; Chan et al. 2021).

While we cannot predict which of the 50 *Perkinsus* tRNAs are imported into the mitochondrion, we examined all for possible explanations for the frameshifts and alternative codons used in *Perkinsus* mitochondria. We identified two tryptophan-like suppressor tRNAs with a TCA anticodon in the nuclear genome of both *P. marinus* and *P. olseni* (Supplementary File 3). These tRNAs are predicted to rescue mtDNA-encoded TGA stop codons providing an alternative anticodon that codes for tryptophan. We also examined tRNAs for potential anticodons complementary to the non-canonical 4-5 bp codes. To this end, we investigated all previously identified RNAs with putative CCA ends from LOTTE-seq for any resemblance to known tRNAs by carrying out BLASTn searches against a database comprising all currently known tRNAs and ncRNAs. No candidates were found, suggesting that the frameshifts in the *Perkinsus* mitochondria are not translated by specialized tRNAs with novel, extended anticodons.

### Unused codons likely trigger frameshift translations

Programmed ribosome frameshifting (PRF) is used by many viruses where it is most often facilitated by two sequence elements: a heptanucleotide ribosomal slippery sequence (or slip-site) followed by a downstream RNA structure (Penn et al. 2020). The slip-site typically has the form X XXY YYZ (X is three identical residues, Y is A or U, and Z is A, U or C). In *Perkinsus*, however, the frameshifting sites to not conform to these slippery sequence motifs (Figure 3A, E). In viruses, RNA secondary structures — either stem-loops or pseudoknots — occur ∼5-10 nucleotides downstream of the frameshift. These features induce kinetic and conformational changes to the ribosome during translation and promote the slipping to the alternative frame (Penn et al. 2020). However, we also found no evidence of these predicted secondary structures in relation to the frameshift sites in *Perkinsus*.

Another mechanism that can contribute to PRF is the relative abundance of tRNAs where relatively depleted tRNAs can promote ribosome stalling and slippage. To test for evidence of such a mechanism in *Perkinsus* we calculated the mitochondrial codon usage frequencies for each *Perkinsus* species. We used the three genes’ predicted reading frames based on the protein alignments (Figures S6-8), only excluding the frameshift sequences from these calculations (AGGY, TMGGY, CCCCT, TAAC) (Figure 3E, Supplementary Table S3). We then plotted the codon-usage frequencies onto the predicted reading frames before and after each frameshift for all genes to assess translatability through these regions. From this analysis, a remarkably conspicuous observation common to all frameshifts was made. At the point of each frameshift a codon that is otherwise unused occurs in the current translation frame, and this is often followed by further unused codons in this frame (Figure 4). In the case of the +1 frameshifts, the first codon of the next frame, and those that follow, are commonly used codons restoring translatability (e.g., COX1: G74). For the +2 frameshifts, the first codon of the next reading frame is also an unused codon, and only after sliding two nucleotides do codons used elsewhere occur again, restoring translatability (e.g., COX1:P80). This situation is seen for all frameshifted positions in all four species (Figure S9-12). These data imply that an absence of select amino acid-charged tRNAs in the mitochondrion results in translation arrest when their cognate codons occur. Translation can only be restored by the entry of an available tRNA that can bind to the next available cognate codon. For COB:G41 in three *Perkinsus* spp., the first frameshifted codon is the TAG stop codon (it is an unused serine codon in *P. chesapeaki*) which is also otherwise unused in the mitochondrial genomes and likely lacks the corresponding release factor, which would again result in ribosome stalling. This mechanism for frameshift translation also provides a prediction for the location of the last frameshift in COB. Translatability is maintained up to the TAA codon at this site, and single nucleotide slippage allows ongoing translation in the final frame starting with an asparagine (Figure 4: COB:N303). Although the translation termination mechanism remains unclear in *Perkinsus* mitochondria, we predict that a delay (or absence) of recognition of this TAA codon allows frameshifting as for the other sites. Protein translatability also implies that the translated protein termini are confined to the highly conserved nucleotide shared between the four taxa (Figure 2). Immediately outside of these regions unused codons are encountered in all three reading frames (Figures S13-14).

**Figure 4.**
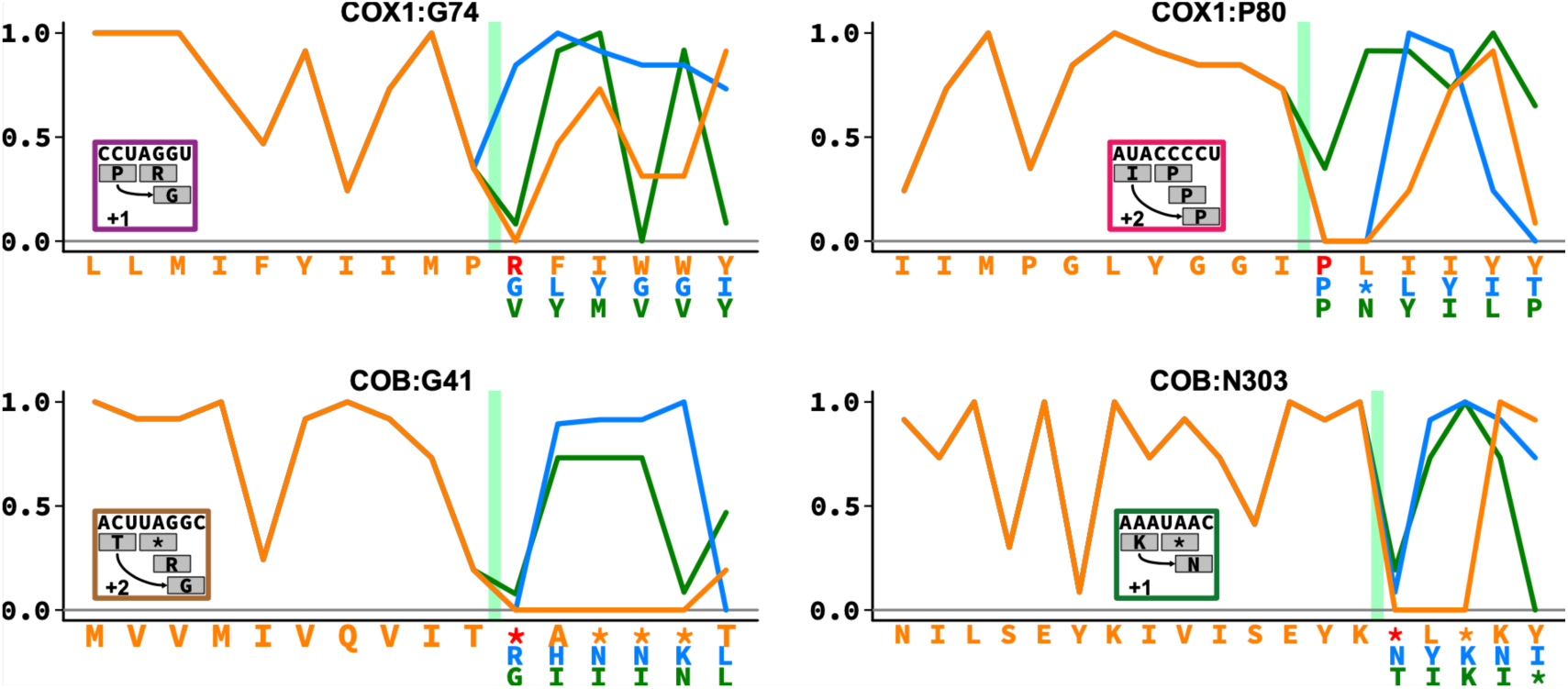
Unused codons coincide with frameshifts. Codon usage plotted for alternative reading frames at representative frameshift sites (green bar: for plots of all frameshift sites see Supplemental Figures S9-12). When the non-shifted sequences (orange) encounter unused or stop codons, the frameshifted sequences (blue: +1 nt frameshift, green: +2 nt frameshift) rescue the translatability of the reading frames. In the cases of COX1:P80 and COB:G41, a +2 frameshift is required because the +1 frameshift also encounters and unused or stop codon.

### Conservation of frameshift sites corresponds to protein secondary structure

The frameshifts in *Perkinsus* spp. mitochondrial genes are predicted to translate as 16 glycines (15 in *P. chesapeaki*), two prolines and one asparagine, however, these amino acids are also coded for by canonical three-nucleotide codons in all three protein-coding genes (Figures S5-7). This raises the question of why the non-canonical codes are used and, given the conservation of their locations in *Perkinsus* spp., if they might perform a position-specific function in their respective proteins. To explore this question, we initially examined the conservation of these glycine, proline and asparagine residues in near and distant orthologues. Alignments of *Perkinsus* COX1 and COB with orthologs shows that frameshifted and canonically coded residues both occur at sites of conserved usage and usage specific to *Perkinsus* (Figure S5-6). For COX3 the site of the single frameshifted glycine does not correspond to a conserved glycine position in other homologues (Figure S7). Moreover, several widely conserved positions for glycine and proline in COX3 lack these residues in *Perkinsus* spp. Thus, for all proteins there does not appear to be a link between protein sequence conservation and position of the frameshifts.

The location of the frameshiftted residues could contribute to the three-dimensional (3D) properties of the proteins, so we modelled the 3D structure of the *P. marinus* proteins and compared them to known structures from the model *Bos taurus*. All structure predictions were of high confidence (c-score >0) and showed near identical matches with the *B. taurus* protein structures (root-mean-square deviations <2) (Figure 5A). The strong fit of the *Perkinsus* proteins further substantiates our identification of *Perkinsus cox3*s and the 3’ extension of *cob* coding sequences beyond the TAA frameshift. The locations of the frameshifted residues showed no obvious pattern with respect to the protein structures: they occurred on both sides of the inner mitochondrial membrane that these proteins span, and both within and between predicted transmembrane helices. However, given that the frameshifts likely cause temporary interruption to translation we speculate that they might contribute to some properties of protein folding. Nascent polypeptides can commence folding into secondary and tertiary structures once they emerge from the ribosome exit tunnel which is typically the equivalent of ∼28 amino acids of the nascent chain. When the region situated 25-30 residues upstream of each frameshift was plotted on the predicted protein secondary structures, this emergent region typically occurred at the N-terminal boundaries of alpha helices (Figure 5B). These observations suggest a pause in translation occurs before the emergence, folding and/or membrane insertion of helical secondary structures. An interesting exception is for COX3 where the single frameshift occurs less than 20 residues from the predicted protein start, suggesting that this translation pause occurs before the N-terminus emerges from the ribosome.

**Figure 5.**
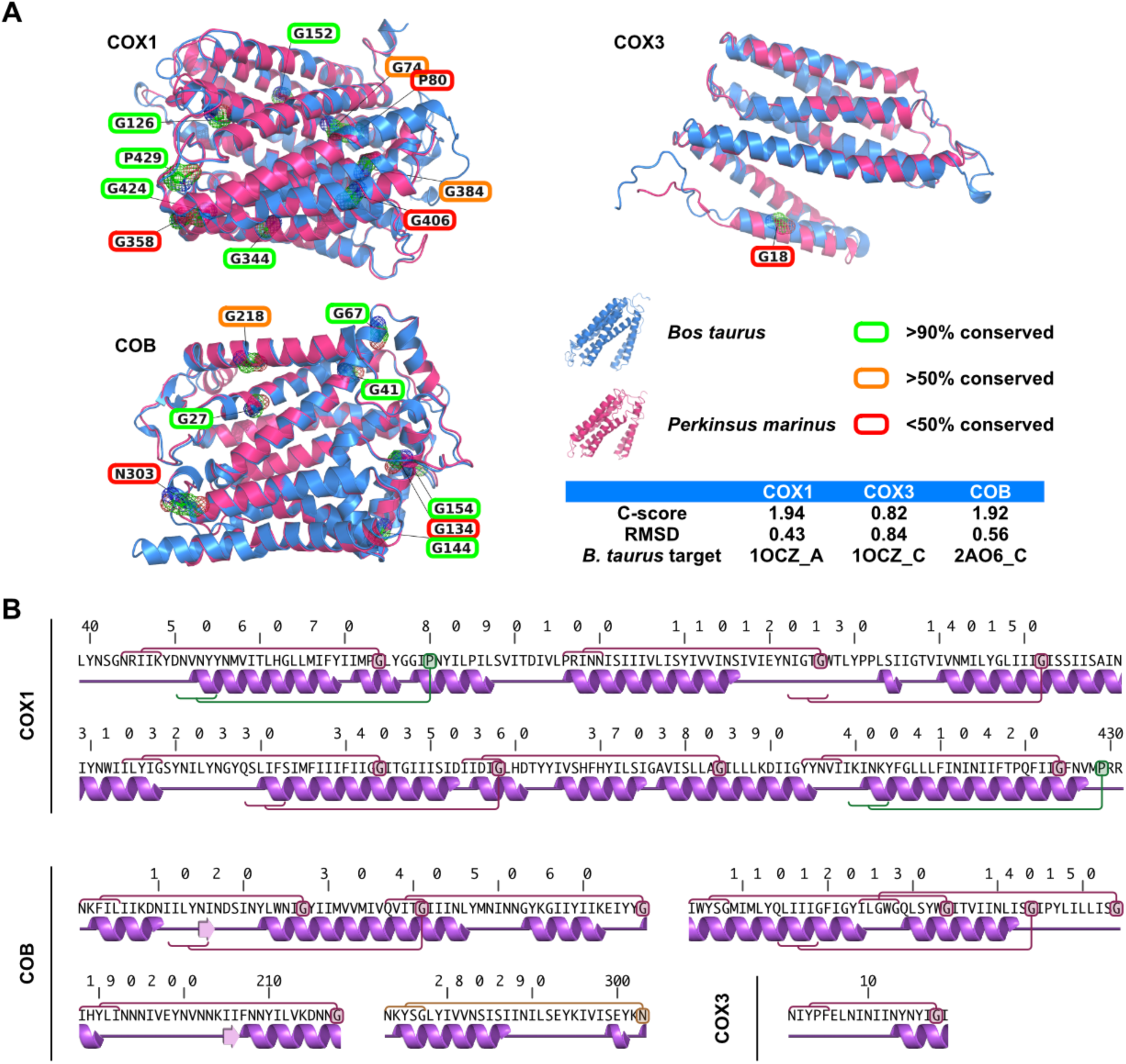
Frameshift position conservation and location with respect to protein structures. ***A***. Frameshifted glycine, asparagine and proline residues (depicted as mesh volumes) in the predicted structures of COX1, COB and COX3 from *P. marinus* (pink) overlaid on *Bos taurus* structures (blue, Protein Data Bank structures: 1OCZ:A, 1OCZ:C and 2A06:C). The non-canonically coded residues are indicated with colored boxes corresponding to their overall conservation (see key and Figure S6-8). Modelling and alignment scores for each protein are shown. ***B***. Predicted secondary structures of COX1, COB and COX3 showing frameshifts (circles) and corresponding predicted protein emergence sites from the ribosome exit tunnel when each frameshift is translated (red brackets). RMSD, root-mean-square deviation.

### Fragmented but incomplete ribosomal RNAs in *Perkinsus* mtDNA

A further feature of myzozoan mtDNAs is the absence of full-length rRNA sequences. In *Plasmodium, Hematodinium* and *Chromera* the presence of heavily fragmented rRNAs has been reported (Feagin et al. 2012; Jackson et al. 2012; Flegontov et al. 2015). No complete rRNAs were found in the *Perkinsus* mtDNAs, so we searched for corresponding fragments to those found in other myzozoans. Using 39 rRNA fragments from *Plasmodium falciparum*, 17 from *Hematodinium* and 7 from *Chromera* as queries, we identified a total of 23 rRNA fragments across the mtDNAs of the four *Perkinsus* species. This is consistent with the rRNA fragmentation having occurred before the divergence of Myzozoa. The representation of rRNA fragments across the four *Perkinsus* taxa, however, was incomplete and not always homogeneous. Only the large ribosomal subunit F fragment (LSUF; as per *P. falciparum* annotation (Russell and Beckenbach 2008; Dinman 2012; Feagin et al. 2012; Seligmann 2012; Haen et al. 2014)) was found in all four mtDNAs (Figure 1, 6 and S15). Other fragments were found in three or less taxa; for example LSUG, LSUC and RNA1. Furthermore, several rRNA fragments conserved in other myzozoans were not detected in *Perkinsus* spp. (Figure 6). In total, eight rRNA fragments were found in the mtDNA for *P. atlanticus*, seven in *P. marinus*, and only four for *P. chesapeaki* and *P. olseni*. To test if the non-uniform detection of rRNA fragments in *Perkinsus* spp. was due to cryptic undetected mtDNA molecules that had not been assembled, we searched all raw reads from the total cell DNA sequence data with the high AT-content corresponding to the mtDNA. No additional rRNA fragments were found in these data suggesting that our mtDNA assembles are complete.

**Figure 6.**
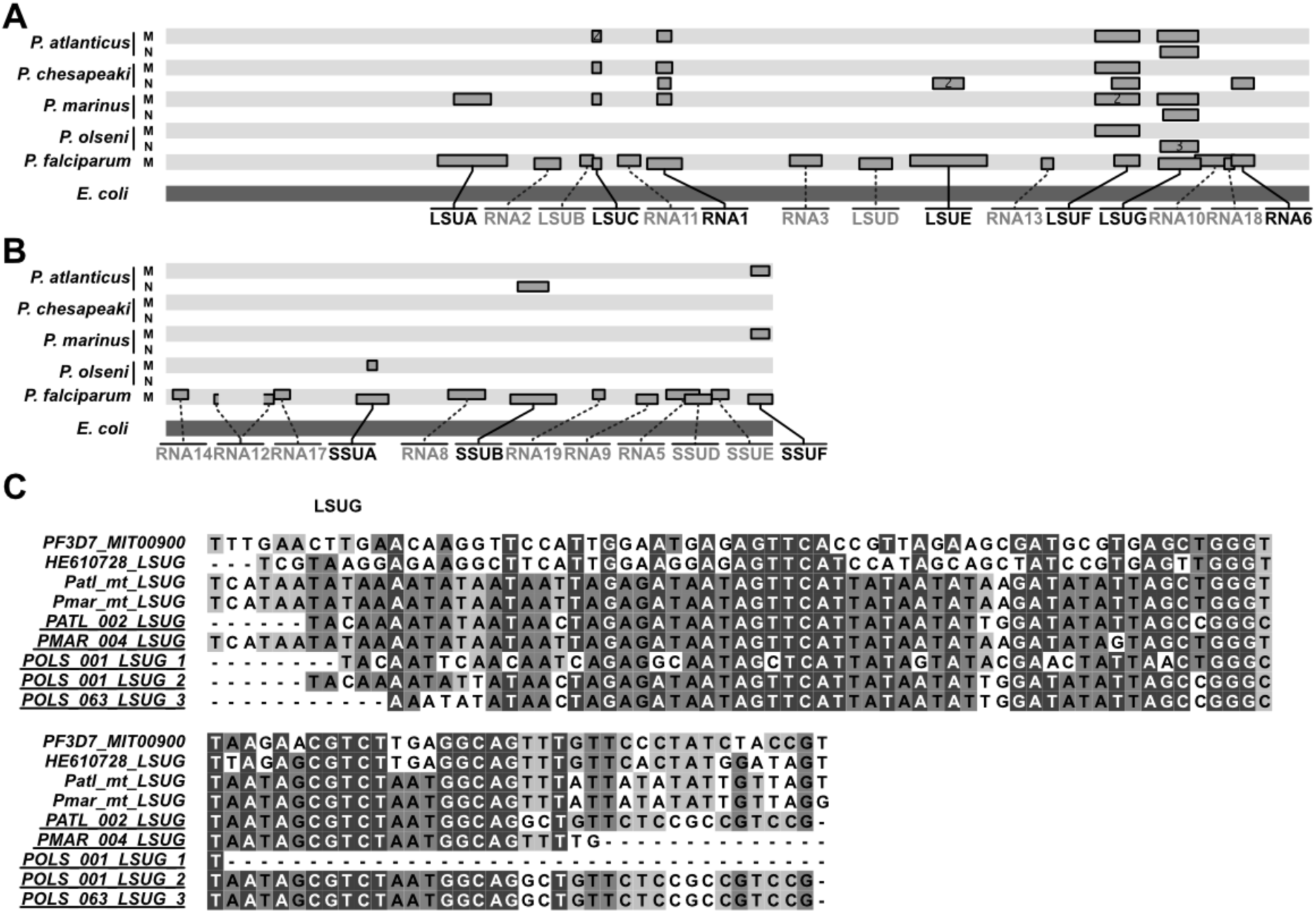
Non-uniform presence of fragmented rRNA sequences in mtDNA and nuclear assembles. ***A***. and ***B***. Distribution of the predicted rRNA fragments of *Perkinsus* with respect to the *Plasmodium falciparum* mt-rRNA fragments and the full-length *E. coli* rRNA subunits (A: large subunit, B: small subunit). The presence and genomic location(s) of the fragments is shown for each *Perkinsus* species (M: mitochondrial; N: nuclear). The correspondence of the *Perkinsus* and *Plasmodium* fragments with respect to the *E. Coli* rRNAs were determined using nhmmer and adjusted according to the structural overlaps of the *Plasmodium* rRNA fragments with respect to canonical rRNA structures (Feagin *et al*., 2012). ***C***. LSUG sequences, as rRNA fragment examples, derived from either mitochondrial (mt) or nuclear genome assemblies (underlined). See Supplemental Figure S15 for further rRNA alignments.

A remaining possible location for mitochondrial rRNAs could be the nucleus. Similar to the mitochondrial tRNAs, it is conceivable that small rRNA fragments would also be amenable to import. We have assembled the nuclear genomes of the four *Perkinsus* taxa from the long- and short-read data (manuscript in prep.) and we searched these assembles also, masking the eukaryotic intact rRNAs. Six putative rRNA fragments were detected within these nuclear assembles (Figure 6, S15). Some fragments were found in both the mitochondrion and nucleus for some taxa (e.g. LSUG for *P. marinus* and *P. atlanticus*) which might indicate recent mtDNA transfer. But for some taxa only a nuclear version of the rRNA fragment was found (e.g. LSUG for *P. olseni*) (Figure 6C). Moreover, some fragments were only found in nuclear assemblies and not in any of the mtDNAs (e.g. SSUB). These data present the intriguing hypothesis that mitochondrial rRNA gene relocation to the nucleus is occurring in *Perkinsus* spp with their transcripts reimported into the organelle.

## DISCUSSION

The myzozoan mtDNAs have unprecedented levels of genome reduction, rearrangement and reconfiguration. Furthermore, they have adopted multiple post-transcriptional processes to restore functional molecules from degenerate genes encoding either proteins or rRNAs. The Perkinsozoa (including the genus *Perkinsus*) were the one major lineage amongst Myzozoa for which the mtDNAs had been largely uncharacterized. Our mtDNA sequences of four *Perkinsus* species have resolved the common ancestral features of myzozoan mtDNA, but they also reveal yet another level of elaboration of the management of mitochondrial encoded genes.

All *Perkinsus* spp. mtDNAs are relatively simple molecules encoding single copies of genes for the three common myzozoan mitochondrial proteins (COB, COX1, COX3) and fragmented rRNAs, and they lack encoded tRNAs. Thus, while the coding capacity is equivalent to other myzozoans, the high level of mtDNA amplification, fragmentation and recombination seen in dinoflagellates, chromerids and select apicomplexans such as *Toxoplasma*, suggests that these latter traits are derived and have occurred multiple times independently (Jackson et al. 2007; Nash et al. 2008; Waller and Jackson 2009; Feagin et al. 2012; Jackson et al. 2012; Flegontov et al. 2015; Berná et al. 2021; Mathur et al. 2021; Namasivayam et al. 2021). Nevertheless, genome rearrangement is seen also in *Perkinsus* spp. with little common synteny or shared intergenic sequence between the four species, and the *P. chesapeaki* mtDNA occurs in two separate molecules. This variability perhaps indicates a general predisposition for myzozoan mtDNA complexity where such states can be tolerated. The apparent use of alternative initiator codons to AUG is also shared throughout Myzozoa (and ciliates also). However, the dinozoan use of mitochondrial RNA-editing and trans-splicing *cox3* mRNAs are evidently derived characters of dinoflagellates only.

*Perkinsus*, however, has also independently gained its own mitochondrial oddities that are otherwise absent from any known myzozoan mtDNAs. The canonical TGA stop codon has been recoded to specify tryptophan, as has occurred independently in ciliates also, and we find candidate suppressor tRNAs that likely enable this. More radical is the adoption of Programmed ribosomal frameshifting (PRF) in all genes, requiring 18 to 19 corrections across the encoded proteome. Our sequencing of four divergent *Perkinsus* taxa shows that the number and position of these frameshifts is very stable. The deepest branching taxon from the others, *P. chesapeaki*, is the only one to show a lost, or possible lack of acquisition, of one frameshift (for COX1 glycine:384). This difference represents only a single nucleotide deletion, and the lack of other such ‘simple corrections’ in any other taxa or positions implies strong selection for the maintenance of these frameshift features.

Programmed ribosomal frameshifting plays an important function in many viral genomes where genome compaction can be achieved by overlapping coding sequences occurring in alternative reading frames (Korniy et al. 2019; Penn et al. 2020). Minus one (−1) frameshifts are most common, but -2, +1 and +2 frameshifts also occur. The frameshift signals typically have two elements, a slippery sequence motif followed by a mRNA structure of stem-loop or pseudoknot. Control of the ratios of alternative products made fulfills the virus’s protein stoichiometric needs. In *Perkinsus*, however, we find no evidence at the sites of frameshifting of either of the signals of this PRF mechanism. Nor are their predicted meaningful products of translation in the alternative frames of the up to 10 sites per gene. In eukaryotic mitochondria there are also rare instances of PRF where the relative abundance of tRNAs, particularly depletion of specific tRNAs, is implicated in ribosome pausing that provides the opportunity for translation shippage into alternative frames (Rosengarten et al. 2008; Russell and Beckenbach 2008; Dinman 2012; Seligmann 2012; Haen et al. 2014). This ‘hungry ribosome’ model is implicated in a *cox3* frameshift in the hexactinellid sponge *Aphrocallistes vastus* where a rare tryptophan codon (UGG) is followed in the +1 frame by a common glycine codon (GGA) (Farabaugh 1996; Beckenbach et al. 2005; Rosengarten et al. 2008). Similarly, in mtDNAs *Polyrhachis* ant species a frameshift in one or two positions in *cob* occurs where a rare codon at the ribosome A-site is combined with a weak wobble pairing of the tRNA bound in the P-site and exact Watson-Crick codon-anticodon pairing in the +1 position that promotes translation in the shifted frame. This mechanism is also consistent with known features of programmed translational frameshifting in the yeast TY1 and TY3 retrotransposons and, furthermore, in bacteria ribosomes are known to slide along a given mRNA and resuming translation several base pairs downstream of a rare codon in certain sequence contexts and/or in physiological conditions of tRNA limitation (Farabaugh 1996; Gallant and Lindsley 1998; Beckenbach et al. 2005).

A strong model for *Perkinsus* mitochondrial PRF driven by tRNA availability is illuminated by the four taxa’s genomes and 75 instances of frameshifts. All frameshifts occur at otherwise unused codons for these mitochondria, and in the case of the +2 frameshifts a further unused codon in the first +1 frame position. Even when there is nucleotide variability between *Perkinsus* spp. at the +2 glycine frameshift (TMGGY), both alternatives of the last codon of the current frame (TAG:Stop, or TCG:serine), and the +1 first codon (GGC:arginine, or GGT:arginine) are all unused codons reinforcing the consistency of this pattern (see Figure 3E). We currently do not know which cytosolic tRNAs are imported into *Perkinsus* mitochondria, however, non-use of the frameshift codons in any other regions of the genes strongly implies that charged tRNAs are absent for these select codons in the mitochondrion. This is likely to also be true for cysteine tRNAs as no in-frame cysteine codons occur in any of the genes indicating the surprising reduction to 19 amino acids of this genetic system. This is despite cysteine being present in all three proteins in non-*Perkinsus* taxa, including occurring at relatively conserved positions (Figure S6-8). An implication for PRF relying on absent tRNAs, particularly where up to 10 frameshifts per gene will demand high fidelity recognition, is that it becomes imperative that these frameshift tRNAs are excluded from the mitochondrion. It might be that mechanism for tight import-regulation of permitted tRNAs could also explain the loss of cysteine from this genome where a propensity for tRNA exclusion could have also led to a permissible loss. mRNA secondary structures might also contribute to *Perkinsus* PRF as our modelling of the most abundant glycine +1 frameshifts (AGGY), used to identify the *cox3* frameshift, identified a strong 11-15 nucleotide consensus sequence flanking this frameshift type. While these sequences do not present obvious stem-loop or pseudoknot secondary structures as for viral and some bacterial PRF, it might be that they recruit specific proteins or short binding nucleic acids that assist in translation stalling and promotion of the change of ribosomal reading frame, as for the viral systems (Korniy et al. 2019; Penn et al. 2020).

The properties of *Perkinsus* mitochondrial ribosomes, while poorly understood, might have contributed to the evolution of abundant frameshifts in their genes. Myzozoan ribosomes are known to be unusual with highly fragmented rRNAs in 20 or more pieces. It is not known how these ribosomes assemble. Conserved rRNA domains of other prokaryote-derived ribosomes are apparently missing in most, and inconsistently present in many, myzozoan groups (Feagin et al. 2012; Jackson et al. 2012; Flegontov et al. 2015). The four *Perkinsus* mtDNAs even show significant variability between species and, tantalizingly, we observe potential for some fragments to have relocated to the nucleus from where their transcripts might be reimported as for tRNAs. These processes could have resulted in ribosomes of significantly altered structure that might also impact their properties. This could include more permissive access at the ribosome A-site to the mRNA and exploration of alternative frame codons by tRNAs, or a greater ribosome susceptibility to mechanical stresses that might be asserted by mRNA structures and/or binding molecules should these act as frameshift cues. In any case, the density of frameshifts in *Perkinsus cox1* and *cob*, much greater than either *cox3* or most other eukaryotic instances of frameshifts, demands that correction of frameshifts is highly efficient as even low frequency errors would be compounded over eight or 10 frameshift sites. *Perkinsus*, therefore, presents a conundrum of a seemingly degenerate translational system that must be highly efficient, and one that has not developed in other myzozoan mitochondria.

The absence of cognate tRNAs at frameshifts most likely cause a significant pause in translation. The frameshift positional conservation across *Perkinsus* taxa, and their alignment with the positions of ribosome emergence of the helical domains of the nascent COX1 and COB proteins, suggests that this feature has been adopted to regulate protein folding and/or insertion of the proteins into the inner mitochondrial membrane. All of the frameshifts could be lost by simple one-or two-nucleotide deletions, therefore their maintenance suggests that they represent a gain of function in this system, rather than being a product of constructive neutral evolution (Lukeš et al. 2011). An outstanding question is why *cox3* differs so markedly from *cox1* and *cob*: 1) it contains only one frameshift, 2) this frameshift is sufficiently close to the 5’ end of the mRNA that the nascent polypeptide is likely non-emergent from the ribosome during frameshift translation, and 3) it is otherwise entirely devoid of glycines and prolines despite most COX3 proteins containing 10-15 of these residues. In dinoflagellates, *cox3* is also distinguished from *cox1* and *cob* in that it is transcribed as two mRNAs that require trans-splicing to for a mature message, and it acquires a UAA stop to its messages during polyadenylation whereas *cox1* and *cob* are predicted to translate through the short poly-A tail adding a poly-lysine C-terminus. In *Perkinsus* the maintenance of a conserved encoded TAA stop in *cox3* similarly further distinguishes it from *cox1* and *cob*. These curious features suggest that some aspects of the synthesis, insertion and the C-terminus of COX3 are recalcitrant to the divergent genetic properties prevalent in dinozoa, although currently we do not understand the basis of this. However, the peculiar lack of glycines and prolines in *Perkinsus* COX3 suggests that the frameshift-dependent translation of these residues also impacts the use of canonical glycine and proline codons. The mechanism for this is currently unclear because glycine and proline are coded elsewhere in *cox1* and *cob* using codons that are not used at the frameshift sites. Moreover, the asparagine codon AAT, which is encoded at the last frameshift of *cob*, is otherwise only present in one frameshift but is an abundantly used codon in *cox3* and all genes. The *cox3* gene, thus, tantalizes us with yet unilluminated aspects of the mechanisms and evolutionary significance of the divergent genetic properties of the mitochondria of *Perkinsus*, and indeed of all myzozoans.

## ACKNOWLEDGEMENTS

This work was supported by grants from the Australian Research Council (DP130100572), the Gordon and Betty Moore Foundation (doi:10.37807/GBMF9194), the King Abdullah University of Science and Technology (KAUST; BAS/1/1020-01-01) and the Deutsche Forschungsgemeinschaft (DFG; MO 634/21-1, MO 634/8-2 and INST 268/413-1). We thank Fathia Ben Rached, Sara Mfarrej and Amit K. Subudhi of KAUST for helping with culturing and *Perkinsus* DNA extractions

## METHODS

### Total DNA extraction, sequencing, and assembly of mtDNAs

High molecular weight genomic DNA of *P. marinus, P. olseni* and *P. chesapeaki* was extracted using the Qiagen genomic tip kit for high molecular weight DNA, following the manufacturer’s instructions and the nucleic acid integrity control was performed using a Fragment Analyzer™ (Advanced Analytical Technologies). The extracted genomic DNA was quantified with a Qubit® 2.0 Fluorometer and was used for the preparation of SMRTbell Libraries which were sequenced using a PacBio RSII sequencer and a PacBio Sequel sequencing platform (Pacific Biosciences). Following assembly with canu v1.9 (Koren et al. 2017) we noticed a small number of redundant, low G+C contigs, which represented full-length and partial mtDNAs. In addition we sequenced all four taxa using standard Illumina designe. Here we extracted genomic DNA for the four *Perkinsus* taxa using a DNeasy Blood and Tissue Kit (Qiagen) according to the manufacturer’s instructions. The DNA was quantitated using the Qubit® 2.0 Fluorometer and then sheared on a Covaris E220 (Covaris) to ∼500 bp. The DNA libraries were made using the TruSeq Nano DNA Library Prep kit (Illumina), according to the manufacturers’ instructions. The amplified libraries were stored in -20 °C. The pooled libraries were sequenced in an Illumina HiSeq4000 instrument (2 × 150 bp PE reads) (Illumina), at KAUST Core Lab facility. A PhiX control library was applied to the sequencing run as a base balanced sequence for the calibration of the instrument so that each base type is captured during the entire run. Using Illumina reads we then used the organelle assembler NOVOPlasty (Dierckxsens et al. 2017) with a seed-and-extend algorithm using a previously reported coding sequences of *cox1 and cob* as seed sequences from Zhang et al. (2011) (Zhang et al. 2011) to assemble *Perkinsus* mtDNAs. This yielded four full-length *Perkinsus* mtDNAs. For *P. marinus* and *P. olseni* both assembly methods yielded identical full-length mtDNAs.

### Total RNA extraction and sequencing

Total RNA for the four *Perkinsus* taxa was extracted using Trizol reagent (Invitrogen) following manufacturers’ instructions and quantitated using the Qubit® 2.0 Fluorometer and was used for library preparation. The strand specific RNA libraries were made using the TruSeq Stranded mRNA Sample Prep Kit according to the manufacturer’s instructions. The amplified libraries were sequenced in an Illumina HiSeq4000 instrument (2 × 150 bp PE reads) at KAUST Core Lab facility. A PhiX control library was applied to the sequencing run as a base balanced sequence for the calibration of the instrument so that each base type is captured during the entire run.

### PCR amplification and Sanger-sequencing of *cox1, cox3* and *cob*

In addition to the short-read sequencing, both DNA and cDNA versions of *cox1, cox3* and *cob* were amplified from *P. marinus* and *P. olseni* and Sanger sequenced. Extracted DNA was processed using GenElute™ Mammalian Genomic DNA Miniprep Kit (Sigma), including the optional RNase A digest. For RNA, pellets were rapidly thawed and lysed in >9 volumes of TRI-Reagent (Sigma) and processed using Direct-zol (Zymo), including an on-column DNase digest. To ensure complete removal of residual DNA, 5 μg of the purified RNA was further treated in solution by 5 units of RQ1 DNase in 100 μl of 1x RQ1 buffer (Promega) for 1 h at 37°C, re-extracted by phenol/chloroform and ethanol-precipitated. For each species, 300 ng of total RNA was reverse-transcribed in 20 μl using Transcriptor (Roche) primed with 150 pmol random hexamers following the manufacturer’s protocol. Parallel reactions were set up without reverse transcriptase (RT-) to ensure no genomic DNA contribution to RNA-derived amplicons. 2.5 μl reverse transcription reactions or ∼10 ng genomic DNA were then subjected to PCR amplification by the high-fidelity Phusion polymerase (NEB) in 50 μl reactions using the primer pairs listed in Supplementary Table S1 and the following cycling program: 30 sec denaturation @ 95°C, 30 cycles of [30 sec denaturation @ 95°C, 30 sec annealing @ 50°C, 45 sec extension @ 72°C], final blunting for 1’ @ 72°C. Primers were designed across stretches of sequence identical between *P. marinus* and *P. olseni* genes, with a predicted melting temperature of ∼51°C (using default parameters in Primer3 http://bioinfo.ut.ee/primer3-0.4.0/primer3/). No PCR product was observed in RT-samples or in no-template controls for either the reverse transcription (RT0) or the PCR step (NTC). Purified PCR products were Sanger-sequenced using Cox1-F, Cox3-R or Cob-F for the corresponding amplicons (Supplementary Table S1).

### LOTTE-seq for tRNAs

Total RNA was extracted from *P. marinus* and *P. olseni*, and prepared for LOTTE-Seq as previously described (Erber et al. 2019). Upon sequencing, LOTTE-Seq reads with genome coverage greater than 100X were filtered, clustered and subjected to the tRNA gene predictor tRNAscan-SE 2.0 (Erber et al. 2019; Chan et al. 2021) to either identify typical tRNAs or atypical tRNAs such as trans-spliced and circularized permuted tRNAs; and to classify them accordingly using the default parameter settings as previously described (Edgar 2004; Erber et al. 2019). Sequences not identified as tRNAs with tRNAscan-SE 2.0 were inspected individually for similarities with tRNAs. Remaining unassigned reads were analysed with BLASTn (https://blast.ncbi.nlm.nih.gov/Blast.cgi) for signs of homology with tRNAs or other known ncRNAs. In addition, their secondary structures, computed using RNAfold 2.0 (Lorenz et al. 2011), was investigated for similarities to tRNAs.

### Phylogenetic analysis

Using a complete *P. marinus* 18S small subunit (SSU) rRNA as search bait (AF126013.1) and BLASTn, an alignment was generated of 146 perkinsid and closely related Alveolata taxa including members of the Syndiniales and core dinoflagellates comprising at least 1221 aligned bases for each taxon using the aligner MUSCLE (Edgar 2004; Capella-Gutierrez et al. 2009) at standard settings within Geneious 8.1.9 (Biomatters). The alignment was automatically trimmed using trimAI (Capella-Gutierrez et al. 2009; Nguyen et al. 2015; Minh et al. 2020) and a maximum likelihood tree was calculated using iqtree at standard settings (Rice et al. 2000; Nguyen et al. 2015; Minh et al. 2020). Branch support was calculated using the Ultrafast Bootstrap [UFBoot] algorithm with 1,000 replicates and the Shimodaira–Hasegawa approximate likelihood ratio test [SH-aLRT] allowing ultrafast bootstrap support values to first converged. The full and trimmed alignments, raw consensus tree file and accession numbers are provided in PHYLIP format (Supplementary File 1).

### Further bioinformatic analyses

Genomic G+C content was determined using the *geecee* program from EMBOSS v6.6.0 (Rice et al. 2000; Bernt et al. 2013; Donath et al. 2019). Automated mtDNA annotation was carried out using MITOS2 (Zhang et al. 2000; Bernt et al. 2013; Donath et al. 2019). For confirmation the coding sequences of *cox1* and *cob* were detected using BLASTn v2.11.0 (Zhang et al. 2000; Blum et al. 2021) using the reported sequences of *cox1* and *cob*. To test for *cox3*, we performed BLASTx searches using the whole mtDNA genome sequences against 74,641 *cox3* proteins. Since BLASTx searches did not reveal the coding sequence of *cox3*, we extracted the ORFs longer than 300 bp using the *getorf* program from EMBOSS. We then subjected the deduced amino acid sequences to interproscan searches (Harris 2007; Blum et al. 2021) and selected the ORFs that presented the domains IPR000298 (Cytochrome c oxidase subunit III-like) and IPR035973 (Cytochrome c oxidase subunit III-like superfamily). Synteny analysis was performed by searching for highly similar regions among the four mtDNAs with BLASTn (task: megablast, word-size = 4, e-value less or equal than 1e-3). Only reciprocal matches were kept for visualization. Whole genome alignments of the mtDNAs were carried out within Geneious R10 (Biomatters) using a LASTZ v1.02.00 (Harris 2007; Krzywinski et al. 2009) plugin applying a step length of 30, a seed pattern of ‘12 of 19’, searching both strands, allowing a single transition in a seed hit and applying a HSP threshold of 3,000. Mitochondrial genome visualization was performed using circos v0.69 (Krzywinski et al. 2009; Ma et al. 2014).

To model the most abundant frameshift sequence occurring in *cox1* and *cob*, we aligned the sequences surrounding such frameshifts and constructed a position specific matrix (PSM) with meme v5.3.0 (Grant et al. 2011; Ma et al. 2014) using the option “one occurrence per sequence”. The PSM was employed to search for potential frameshifts in *cox3* using fimo v5.3.0 (Gouy et al. 2010; Grant et al. 2011) with a q-value threshold of 0.0003. An updated PSM was constructed from the 59 single-nucleotide frameshifts detected in *cox1, cob* and *cox3* (consensus: WWWTAWWWWYT**AGGT**WTAWHAKTWAT, e-value: 4×10^−140^).

COX1, COX3 and COB alignments were constructed using clustal omega [Sievers *et al*., 2011], manually edited using seaview v5.0.4 (Gouy et al. 2010; Okonechnikov et al. 2012) and visualized using ugene (v34) (Okonechnikov et al. 2012; Yang and Zhang 2015). The 3D protein structures were modelled using the I-TASSER (Wheeler and Eddy 2013; Yang and Zhang 2015) server and the resulting models were visualized using the PyMOL Molecular Graphics System (Version 2.4.0 Schrödinger, LLC) and pdbsum (Laskowski et al. 2018). Codon usage was calculated for each species using the cusp program from EMBOSS (Rice et al. 2000) and the predicted reading frames of the full CDS but excluding the 4-or 5-nucleotide frameshift sites. Codon usage frequencies were then plotted for the alternative frames surrounding each predicted frameshift position (−10 to +5 codons).

Sequences corresponding to possible rRNA fragments were searched in the mtDNA genomes using *nhmmer* (Masuda et al. 2010; Zhang et al. 2011; Jackson et al. 2012; Wheeler and Eddy 2013; Bogema et al. 2021) with 39 rRNA fragments from *Plasmodium falciparum*, 17 fragments from *Hematodinium* and 7 fragments from *Chromera velia* as queries. The gap-opening and gap-extension probabilities used for constructing the hidden Markov models of *nhmmer* were adjusted to 0.001 and 0.2, respectively. All heuristic searching parameters were turned off to increase sensitivity. 23 putative rRNA fragments were identified in the mitochondrial genomes. The predicted rRNA fragments and the known rRNA fragments were used as *nhmmer* queries to search for additional rRNA fragments in the nuclear genome sequences of the four species (manuscript in preparation). To avoid detection of cytosolic rRNAs, the regions corresponding to known rRNA genes and transposable elements up to 1000 nt long with metazoan-only Dfam rRNA domains were excluded from the searches.

## SUPPLEMENTARY MATERIALS

**Supplementary Figure S1**

Complete 18S phylogenetic tree shown in Figure 1A.

**Supplementary Figure S2**

LASTZ whole mtDNA alignments showing location and orientation of syntenic blocks relative to the reference sequence for each mtDNA molecule and species.

**Supplementary Figure 3**.

Nucleotide alignment of the *cox1* sequences in all four *Perkinsus* species. Conserved asparagine and phenylalanine codons at the 5’ end of the genes are shown (green and magenta boxes, respectively). Canonical TAA stop codons (red boxes) are found several bases downstream of the last conserved codon (blue box)

**Supplementary Figure 4**.

Nucleotide alignment of the *cob* sequences in all four *Perkinsus* species. Conserved asparagine and phenylalanine codons at the 5’ end of the genes are shown (green and magenta boxes, respectively). The sequence similarity is maintained after the skipped in-frame TAA stop codon (orange box). Further, putative stop codons (red boxes) occur several bases downstream of the last conserved codon (blue box).

**Supplementary Figure 5**.

Nucleotide alignment of the *cox3* sequences in all four *Perkinsus* species. Conserved asparagine and phenylalanine codons at the 5’ end of the genes are shown (green and magenta boxes, respectively). The sequence similarity stops shortly after the TAA stop codon (red box).

**Supplementary Figure 6**.

Sequence alignment of COX1 proteins from *Perkinsus* and related organisms. Conservation is represented by the histogram on top of the alignment. The position of the conserved residues is highlighted.

**Supplementary Figure 7**.

Sequence alignment of COB proteins from *Perkinsus* and related organisms. Conservation is represented by the histogram on top of the alignment. The position of the conserved residues is highlighted (see legend). There are several conserved amino acids (black arrow) after the last predicted frameshift (Asn 303).

**Supplementary Figure 8**

Sequence alignment of COX3 from four *Perkinsus* species and related species. Conservation is represented by the histogram on top of the alignment. Colored boxes indicate the positions in which glycine (orange) or proline (blue) residues are conserved in other species but missing in *Perkinsus*.

**Supplementary Figure 9**.

Codon usage at the predicted frameshift sites (green vertical bars) of *Perkinsus atlanticus*. The non-shifted sequences (orange) produce either stop codons or extremely rare codons, whereas the frameshifted (blue: +1 nt frameshift, green: +2 nt frameshift) sequences restore the translatability of the reading frame.

**Supplementary Figure 10**.

Codon usage at the predicted frameshift sites (green vertical bars) of *Perkinsus chesapeaki*. The non-shifted sequences (orange) produce either stop codons or extremely rare codons, whereas the frameshifted (blue: +1 nt frameshift, green: +2 nt frameshift) sequences restore the translatability of the reading frame.

**Supplementary Figure 11**.

Codon usage at the predicted frameshift sites (green vertical bars) of Perkinsus *marinus*. The non-shifted sequences (orange) produce either stop codons or extremely rare codons, whereas the frameshifted (blue: +1 nt frameshift, green: +2 nt frameshift) sequences restore the translatability of the reading frame.

**Supplementary Figure 12**.

Codon usage at the predicted frameshift sites (green vertical bars) of *Perkinsus olseni*. The non-shifted sequences (orange) produce either stop codons or extremely rare codons, whereas the frameshifted (blue: +1 nt frameshift, green: +2 nt frameshift) sequences restore the translatability of the reading frame.

**Supplementary Figure 13**.

Possible translation start sites of the 3’ coding sequences among the four compared species. Conserved residues at the 5’ end (potential translation start sites) are shown in purple boxes

**Supplementary Figure 14**.

Codon usage at the 3’ ends of the coding sequences. The plots show the codon usage for ten codons prior to the last conserved residue (purple boxes) and five codons past the first TAA codon (red boxes).

**Supplementary Figure 15**

Predicted rRNA fragments in the mitochondrial and nuclear genomes (underlined names) of four *Perkinsus* species. rRNA fragments were named according to their similarity to known *Plasmodium falciparum* and *Hematodinium* mt-rRNA fragments (top sequences in each block).

**Supplementary File 1**

Phylogenetic tree raw files

**Supplementary File 2**

mtDNAs_genbank_files

**Supplementary File 3**

Identification of Trp(W)-TCA_tRNAs

**Supplementary Table 1**

Amplification - sequencing primers

**Supplementary Table 2**

LOTTE-seq_tRNA-pool

**Supplementary Table 3**

mtDNA codon usage adjusted CDS

