## Supplementary figures and images for "Mitochondrial genomes in *Perkinsus* decode conserved frameshifts in all genes"

### Supplemental Figure 1

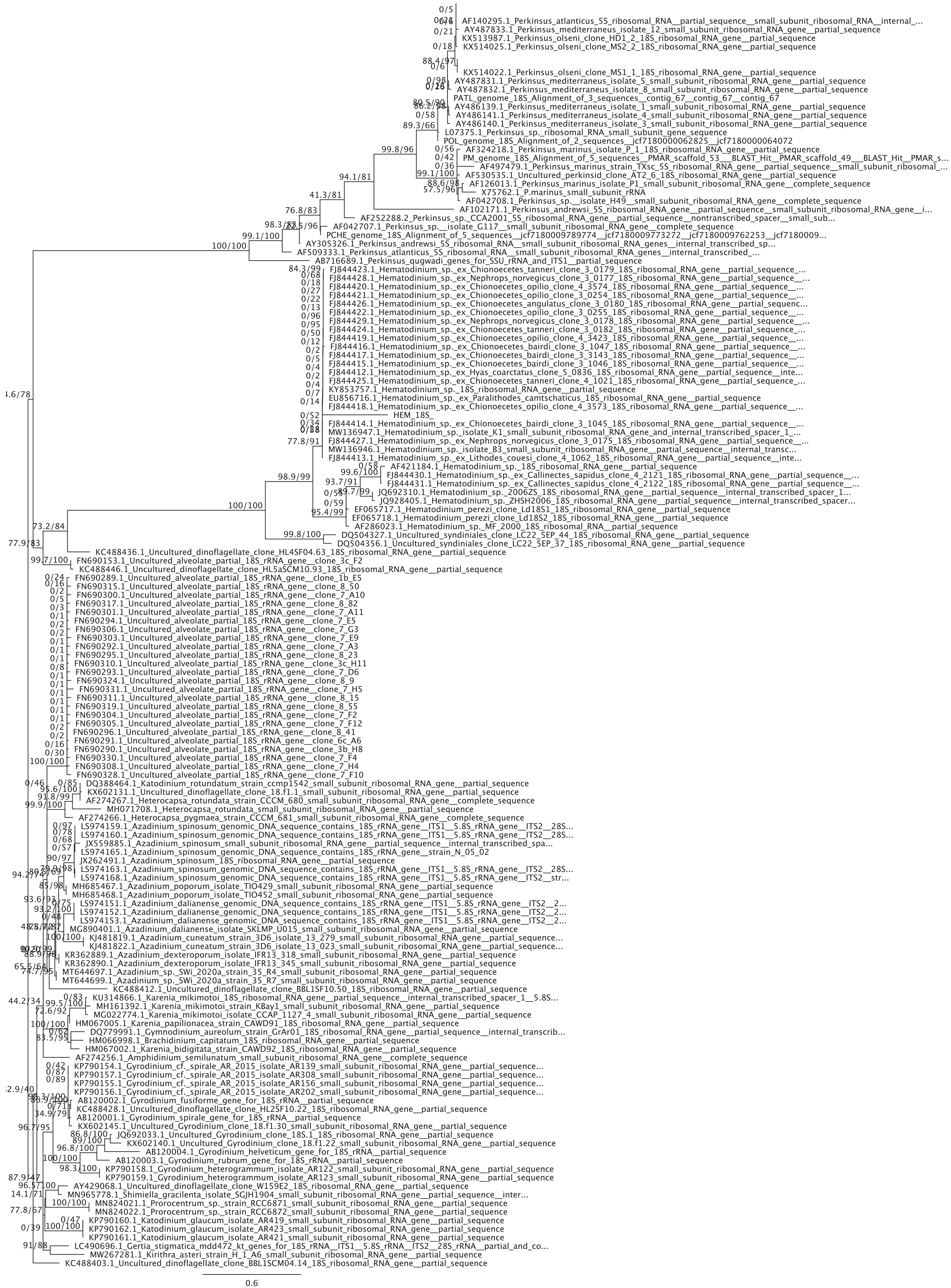

### Supplemental Figure 2

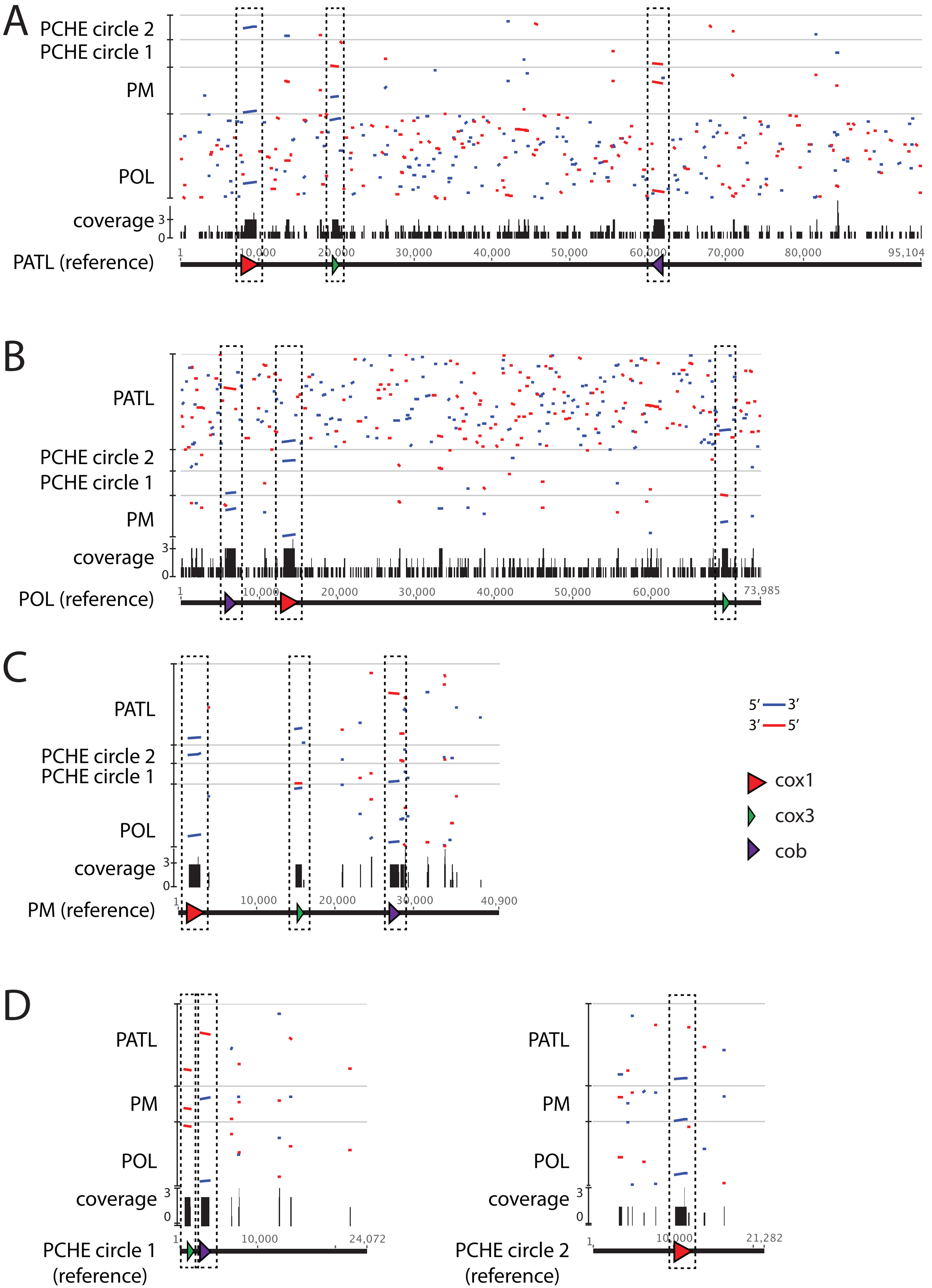

### Supplemental Figure 3

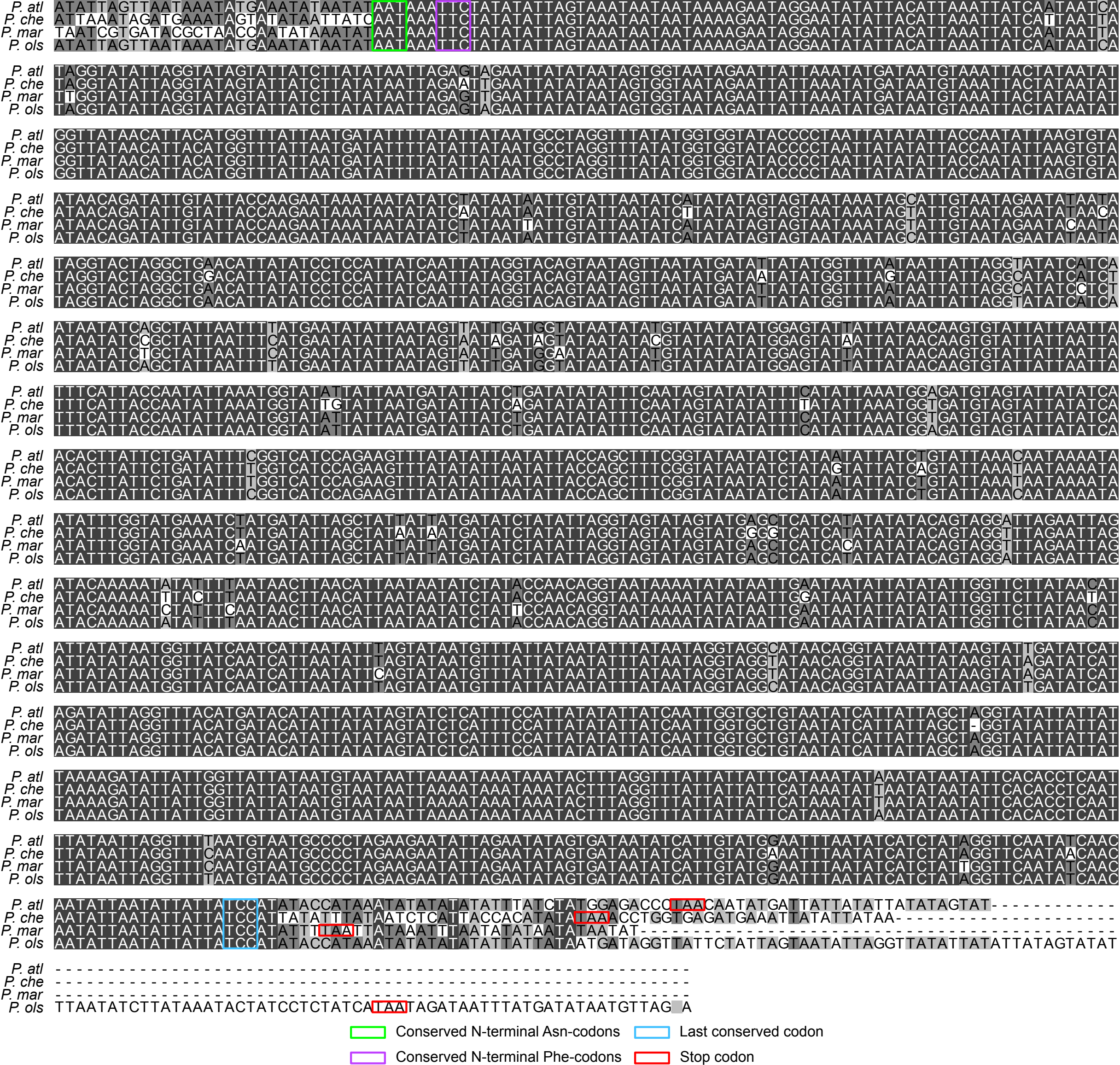

### Supplemental Figure 4

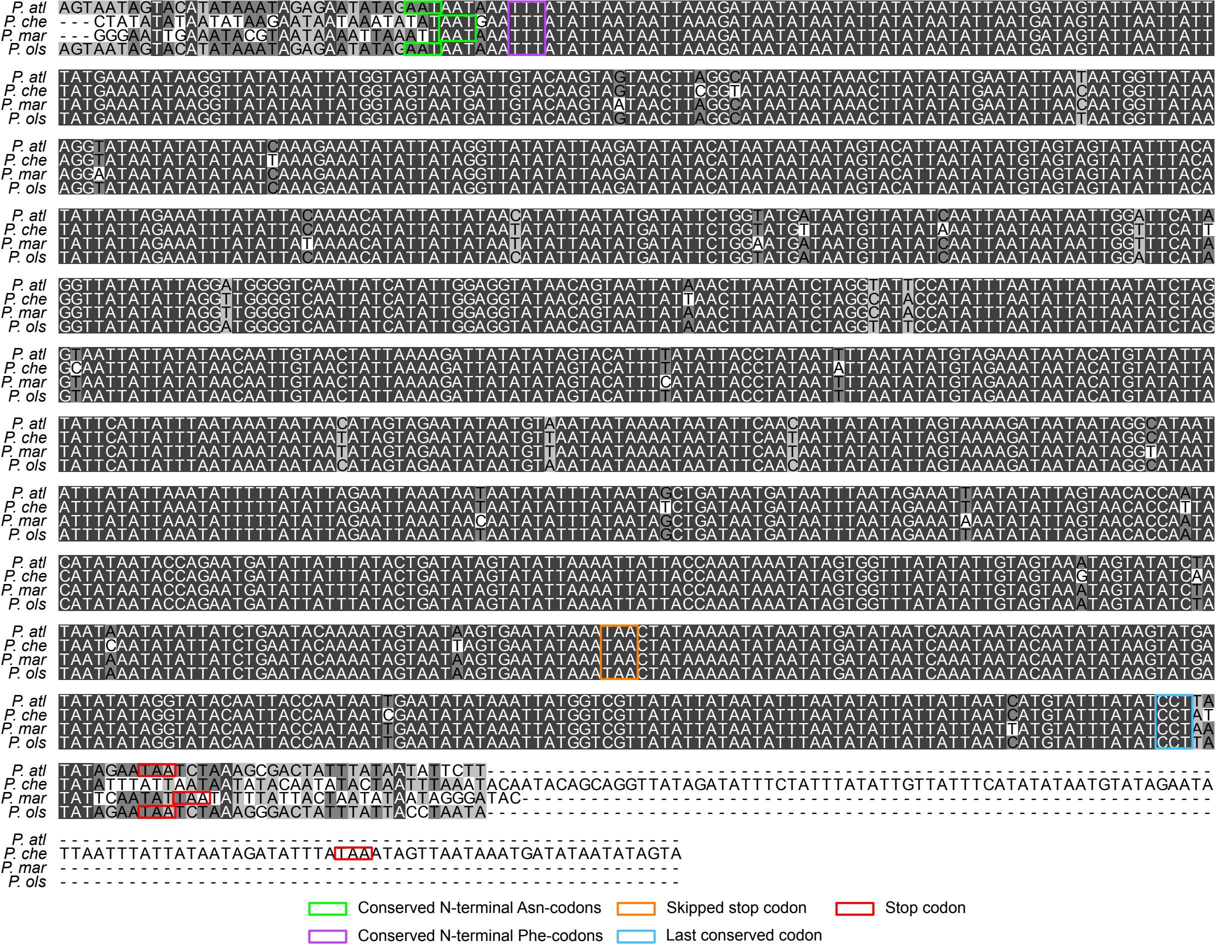

### Supplemental Figure 5

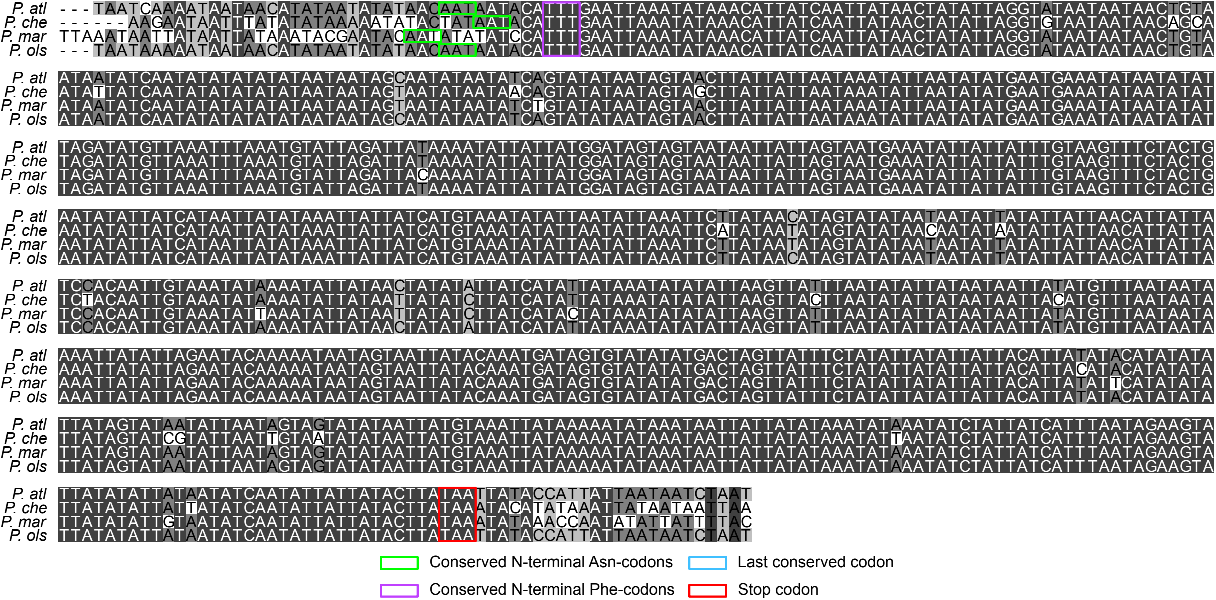

### Supplemental Figure 6

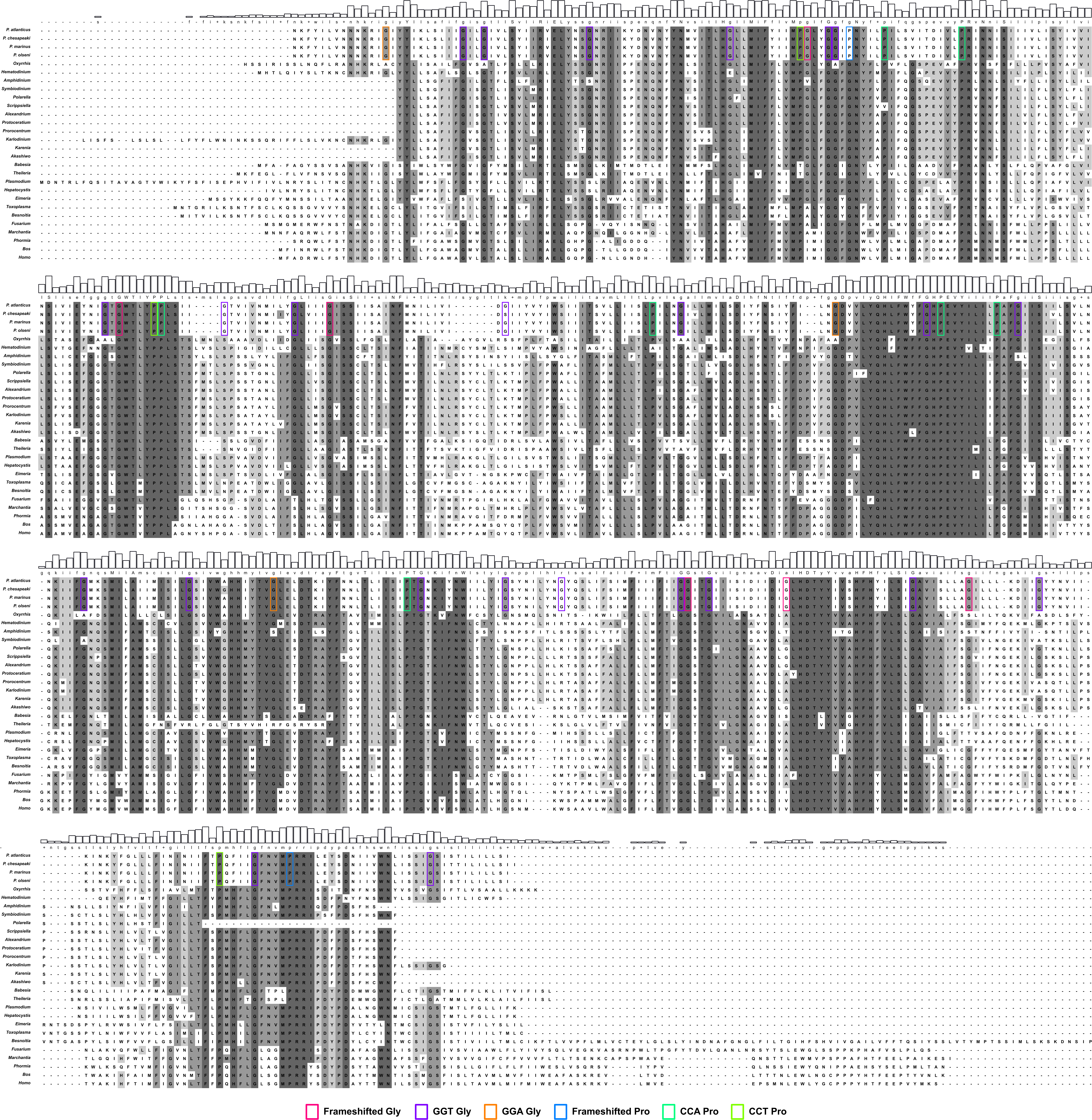

### Supplemental Figure 7

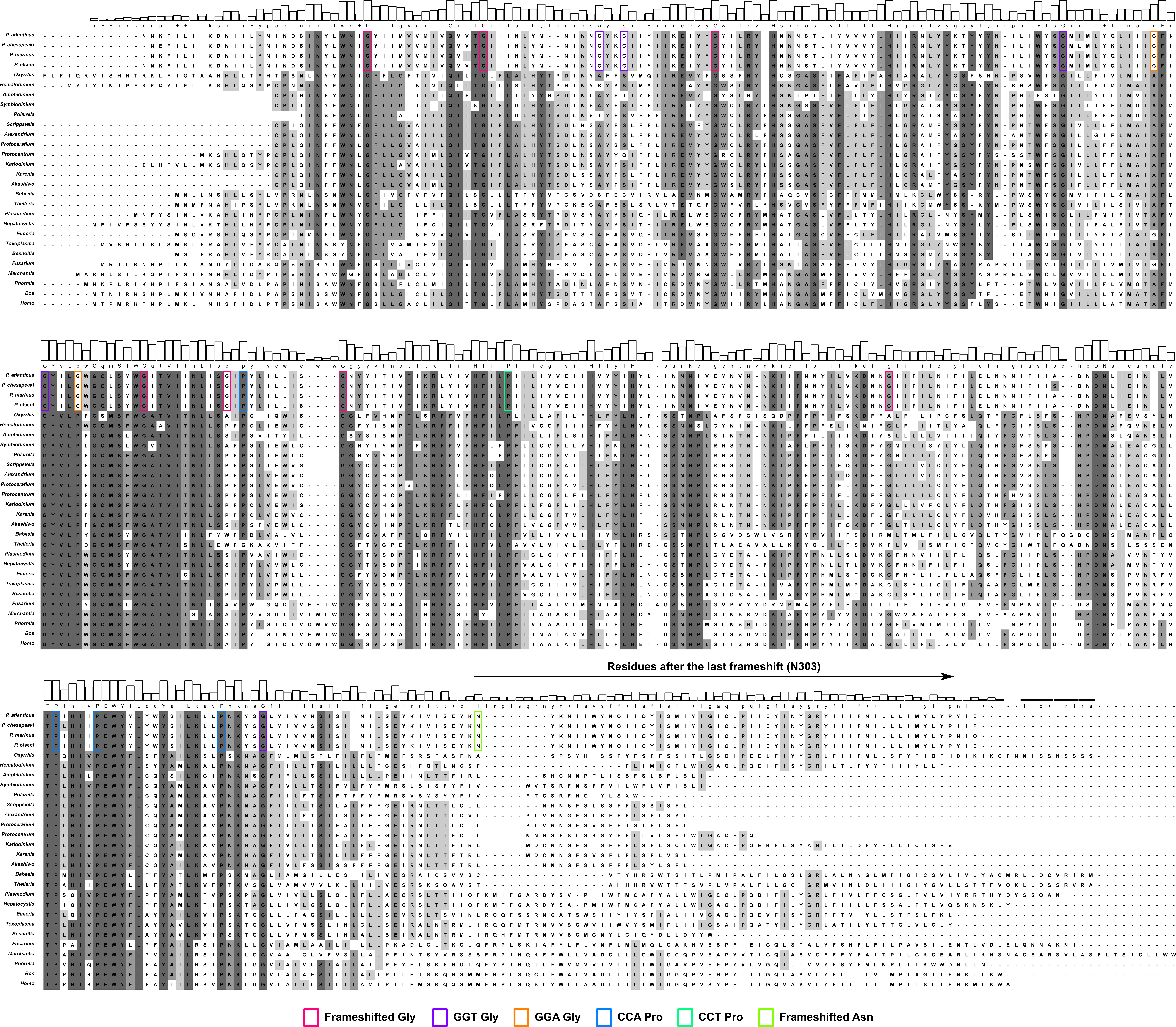

### Supplemental Figure 8

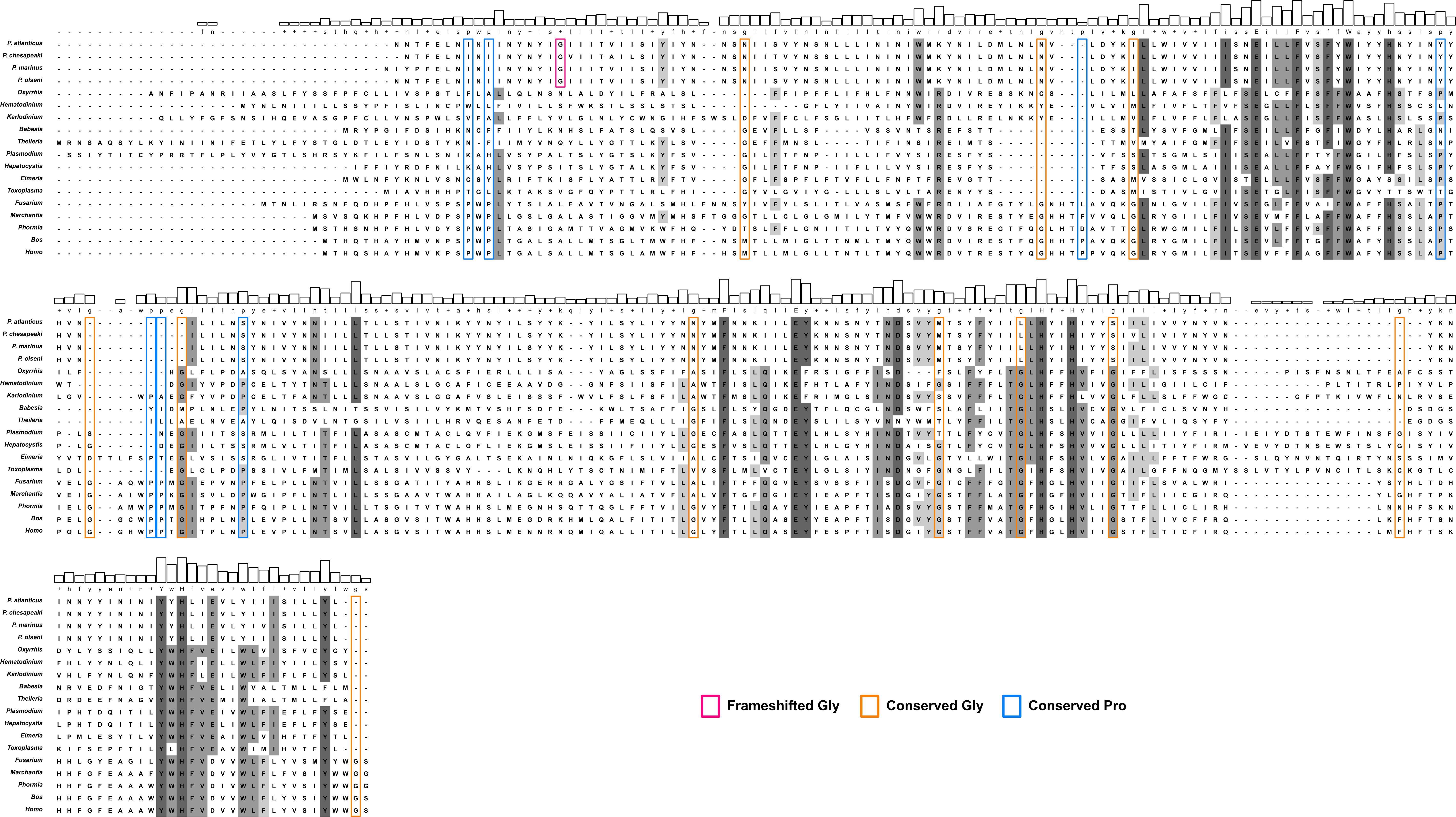

### Supplemental Figure 9

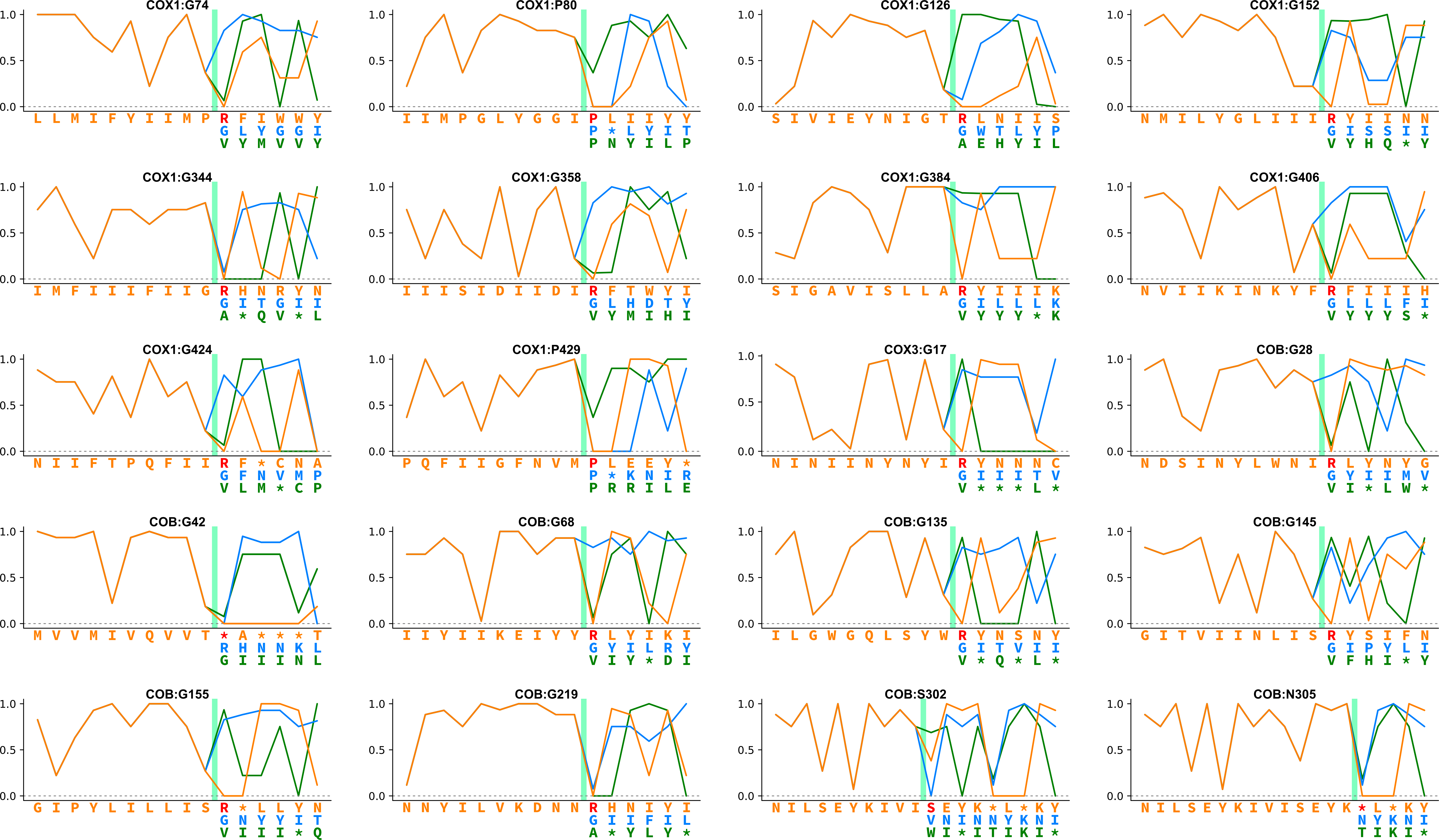

### Supplemental Figure 10

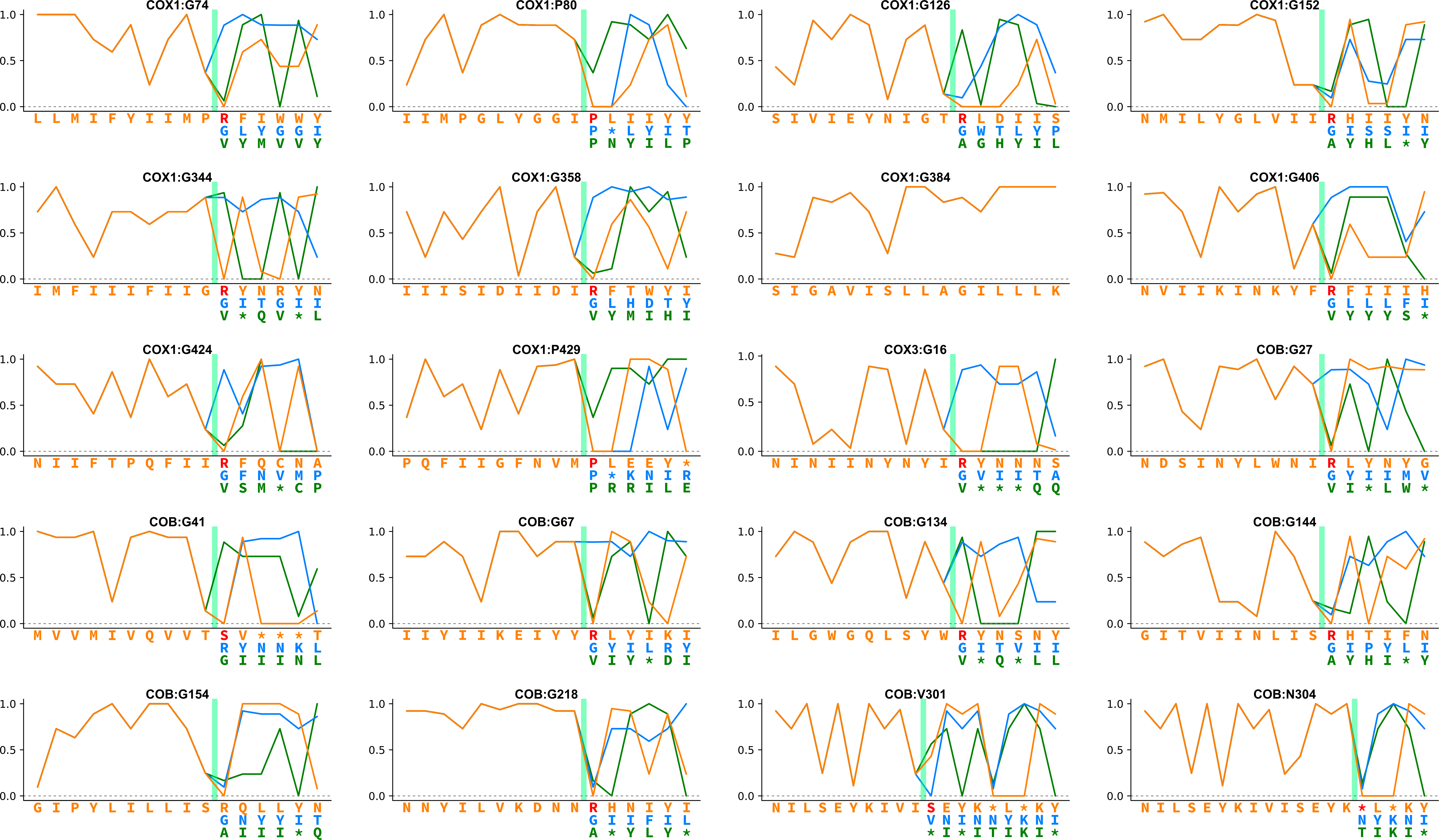

### Supplemental Figure 11

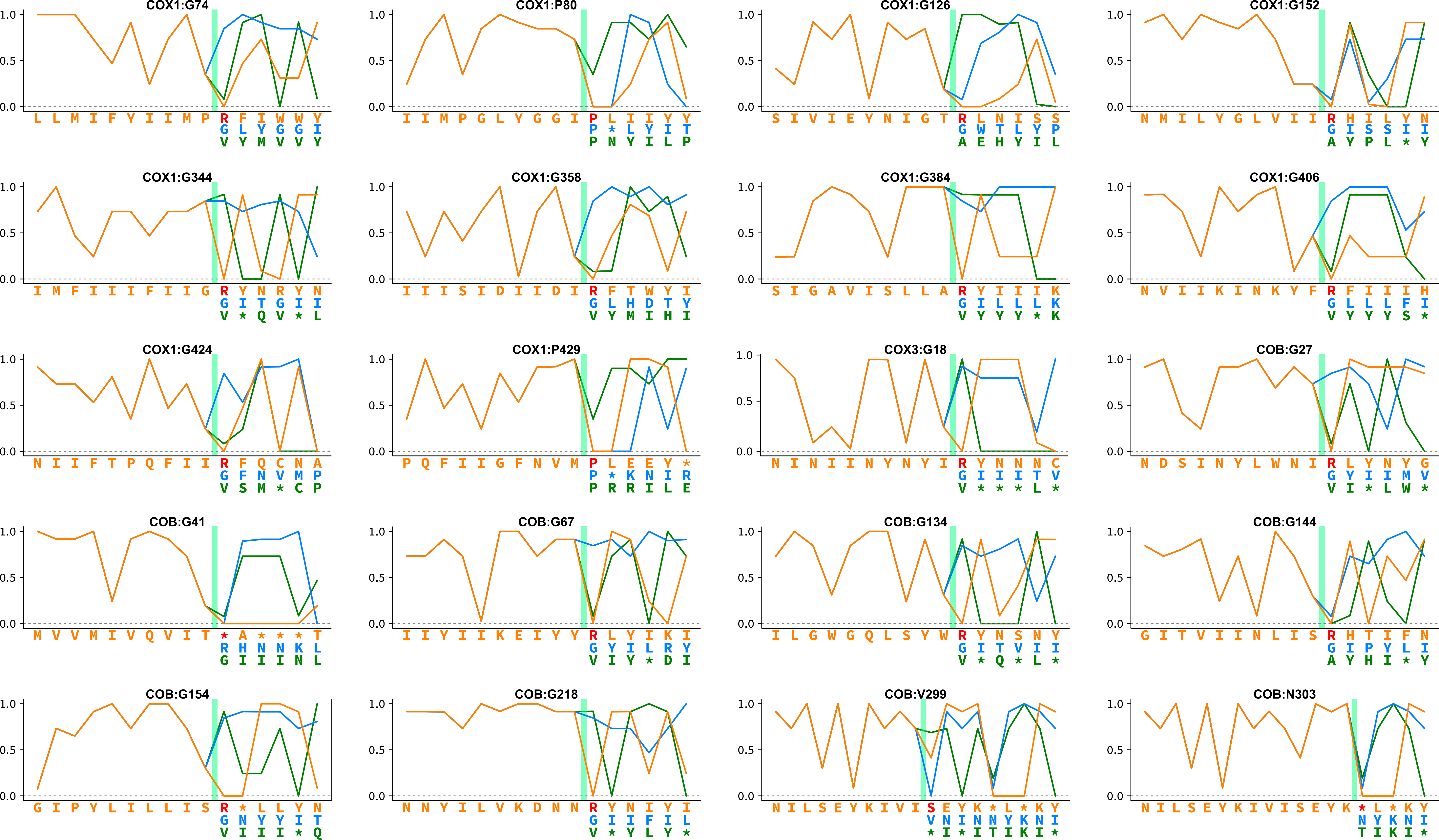

### Supplemental Figure 12

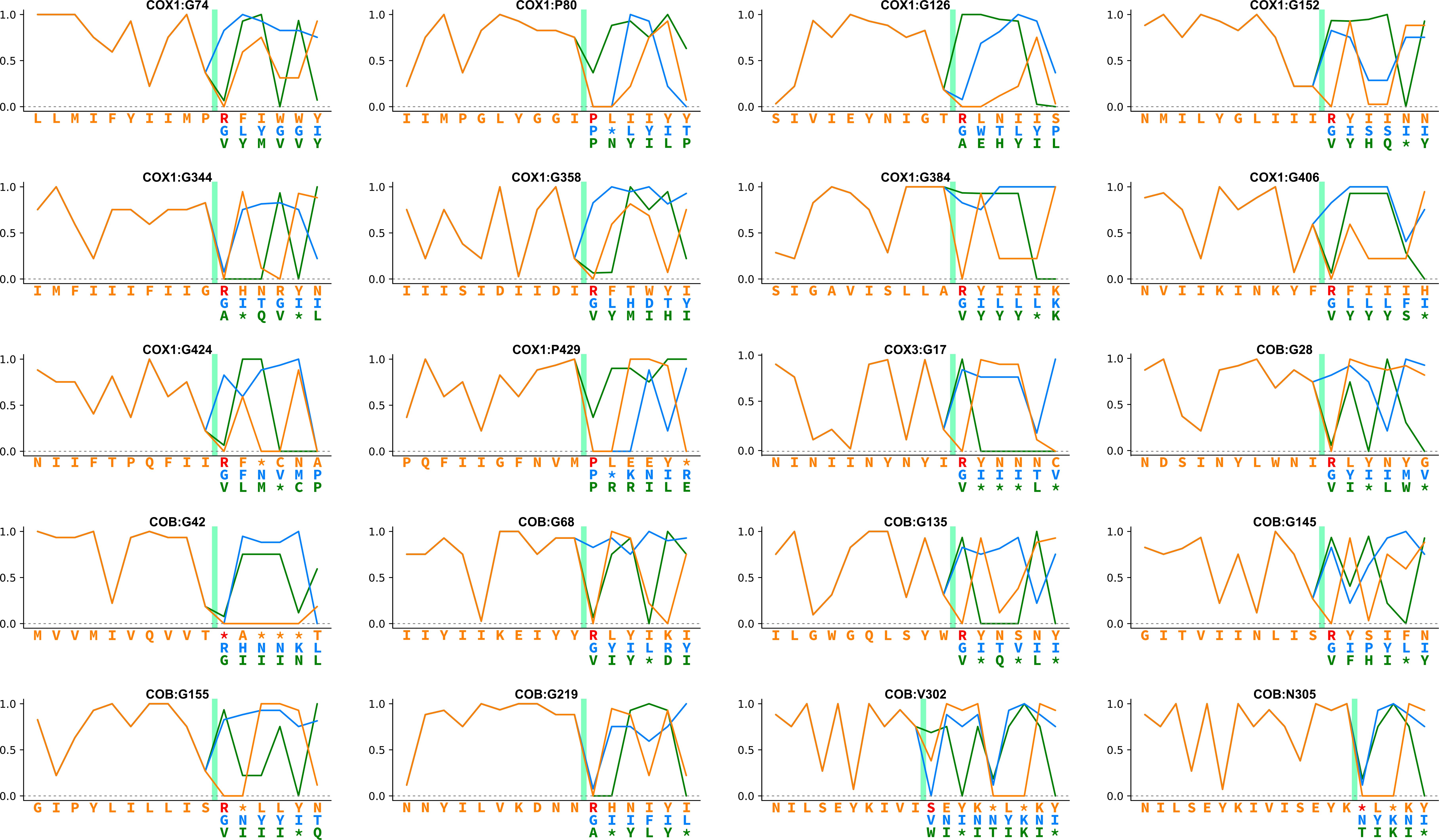

### Supplemental Figure 13

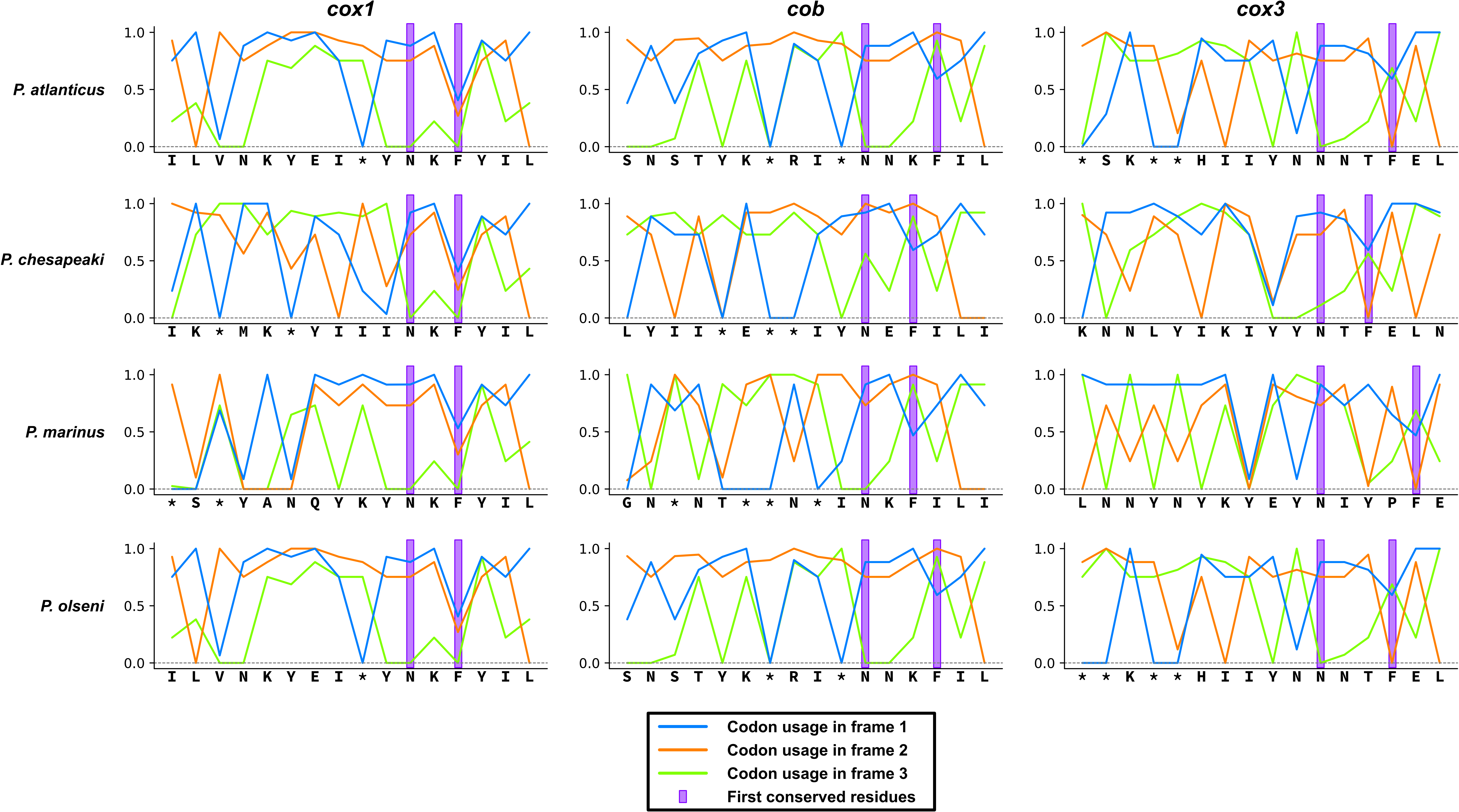

### Supplemental Figure 14

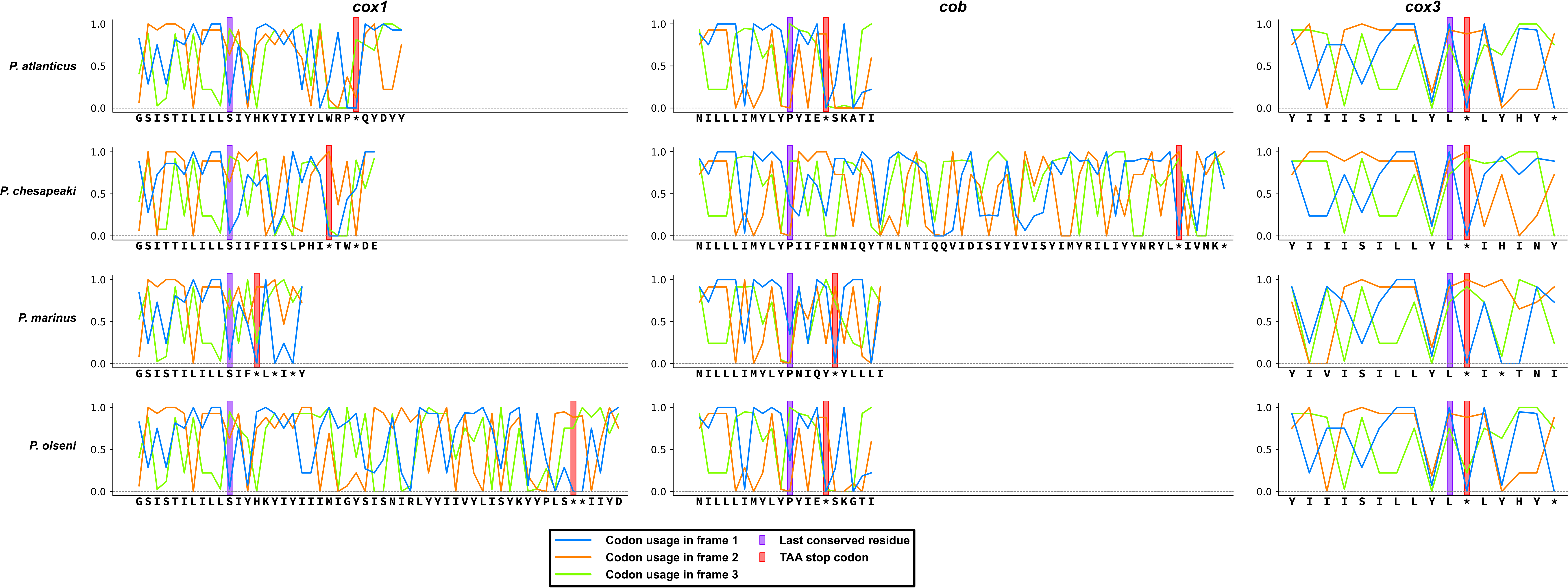

### Supplemental Figure 15

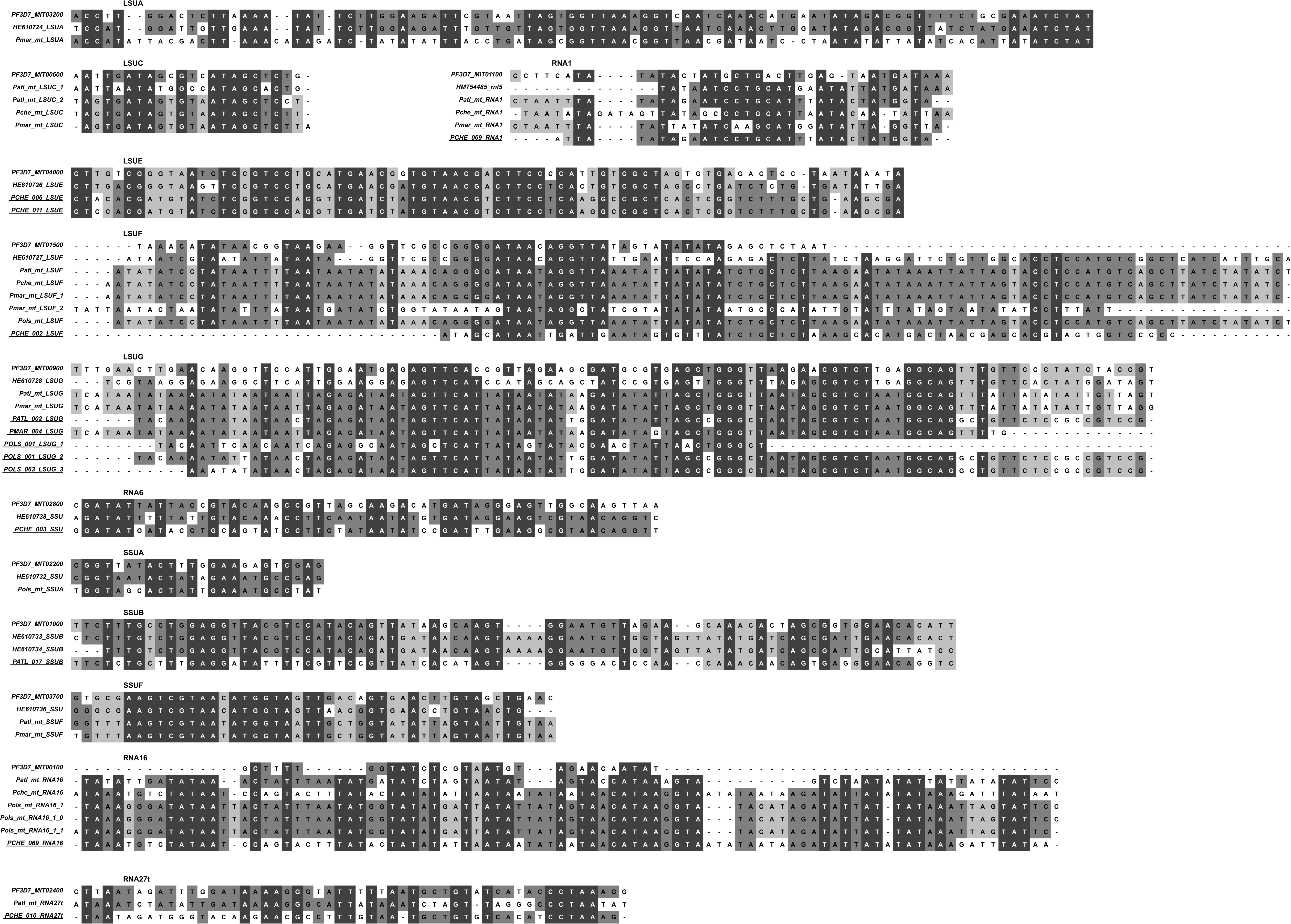
