## Supplemental File 3 for "Mitochondrial genomes in *Perkinsus* decode conserved frameshifts in all genes"

**Identification of alternative Trp-tRNA in *P. marinus* and *P. olseni***

Using (1) tRNAscan-SE2.0 in the nuclear genomes of *P. marinus* (PMAR) and *P. olseni* (POLS) and (2) LOTTE-Seq we identify the following Trp-tRNA like tRNAs with TCA anti-codon (TGA codon).

**(1) *tRNAscan-SE2.0:***

**genomic scaffold name	tRNA begin/end type	anti-codon	intron begin/end	Score**

PMAR_002	1389877	1389806	Sup	TCA	0	0	53.3

PMAR_019	723759	723830	Sup	TCA	0	0	57.0

POLS_001	413668	413597	Sup	TCA	0	0	63.3

POLS_026	363056	363124	Sup	TCA	0	0	37.3 (pseudo)

**(2) *LOTTE-Seq:***

No full-length LOTTE-seq reads corresponding to Trp-tRNA like tRNAs were found in read data, however, shorter reads (below the tRNAscan size threshold) were manual overlapped with loci from tRNAScan results from the genomes. Of these mapped reads secondary structures were considered and allowing identification/confirmation of low coverage alternative Trp-tRNA like tRNAs.

PMAR_002	1389861	1389814

GGTAACGCATCTAACTTCAGATCAGAAGGTTGTCCGTTCGAATCGGGC

PMAR_002	1389873	1389806

CTGTGGCGTAGTGGTAACGCATCTAACTTCAGATCAGAAGGTTGTCCGTTCGAATCGGGCCGGGGTCA

PMAR_019	723759	723830

GGCCCTGTGGCGCAACGGTAGCGCATCTGACTTCAGATCAGAAGGTTGTACGTTCGAATCGGGCCGAGGTCA

POLS_001	413665	413597

CCTGTGGCGCAACGGTAGCGCATCTGACTTCAGATCAGGAGGTTGTCCGTTCGAGTCGGGCCGGGGTCA

POLS_026	363056	363124

GGCCCTGTGGCGCAACGGTAGCGCATTTTTCAGATCAGAAGGTTGTCCGTTCGAATCGGGCCGGGGTCA

Note: in-silico predictions and LOTTE-seq data have good overlap.
